# Diverse molecular mechanisms contribute to differential expression of human duplicated genes

**DOI:** 10.1101/2020.11.27.401752

**Authors:** Colin J. Shew, Paulina Carmona-Mora, Daniela C. Soto, Mira Mastoras, Elizabeth Roberts, Joseph Rosas, Dhriti Jagannathan, Gulhan Kaya, Henriette O’Geene, Megan Y. Dennis

**Affiliations:** Genome Center, University of California, Davis, CA, USA; Integrative Genetics and Genomics Graduate Group, University of California, Davis, CA, USA; MIND Institute, University of California, Davis, CA, USA; Autism Research Training Program, University of California, Davis, CA, USA; Postbaccalaureate Research Education Program, University of California, Davis, CA, USA; Department of Biochemistry & Molecular Medicine, University of California, Davis, CA, USA

**Author notes:** Corresponding author: Megan Y. Dennis, Ph.D., University of California, Davis, School of Medicine, One Shields Avenue, Genome Center, 4303 GBSF, Davis, CA 95616.

## Abstract

Emerging evidence links genes within human-specific segmental duplications (HSDs) to traits and diseases unique to our species. Strikingly, despite being nearly identical by sequence (>98.5%), paralogous HSD genes are differentially expressed across human cell and tissue types, though the underlying mechanisms have not been examined. We compared cross-tissue mRNA levels of 75 HSD genes from 30 families between humans and chimpanzees and found expression patterns consistent with pseudo- or neofunctionalization. In general, ancestral paralogs exhibited greatest expression conservation with chimpanzee orthologs, though exceptions suggest certain derived paralogs may retain or supplant ancestral functions. Concordantly, analysis of long-read isoform sequencing datasets from diverse human tissues and cell lines found that about half of derived paralogs exhibited globally lower expression. To understand mechanisms underlying these differences, we leveraged data from human lymphoblastoid cell lines (LCLs) and found no relationship between paralogous expression divergence and post-transcriptional regulation, sequence divergence, or copy number variation. Considering *cis*-regulation, we reanalyzed ENCODE data and recovered hundreds of previously unidentified candidate CREs in HSDs. We also generated large-insert ChIP-sequencing data for active chromatin features in an LCL to better distinguish paralogous regions. Some duplicated CREs were sufficient to drive differential reporter activity, suggesting they may contribute to divergent *cis*-regulation of paralogous genes. This work provides evidence that *cis*-regulatory divergence contributes to novel expression patterns of recent gene duplicates in humans.

## INTRODUCTION

Gene duplication occurs universally and is considered a major source of evolutionary novelty; across eukaryotes, over 30% of genes are thought to have arisen from duplications (Zhang 2003). Although most duplicated genes rapidly become pseudogenes, some may share and maintain important ancestral functions via subfunctionalization, or gain novel functions entirely (neofunctionalization) (Lynch 2000). Expression divergence is likely integral to the survival of paralogous genes, as spatiotemporal partitioning of function places both daughter paralogs under purifying selection helping them escape pseudogenization (Rodin and Riggs 2003; Rodin et al. 2005). This may be the primary driver of duplicate gene retention, as gene regulation can be altered relatively easily while coding sequences remain intact (Ohno 1970). For example, mouse *Hoxa1* and *Hoxb1* genes are functionally redundant but partitioned by expression, with normal development possible from a single gene under the control of regulatory elements from both paralogs (Tvrdik and Capecchi 2006). On a genome-wide scale, substantial expression divergence has been observed in vertebrates following whole-genome duplications specific to teleost and salmonid fishes (Kassahn et al. 2009; Braasch et al. 2016; Lien et al. 2016; Varadharajan et al. 2018). Meta-analysis suggests that across all these species, selection on gene-expression levels appears relaxed in one of the paralogs (Sandve et al. 2018). However, segmental duplications (SDs, regions defined as having >90% sequence similarity and being at least 1 kb in size (Bailey 2002)) occur more commonly in vertebrates than whole-genome duplications and concomitantly generate structural rearrangements, potentially facilitating regulatory divergence and duplicate retention (Rodin et al. 2005). Although comparative studies characterizing expression divergence of duplicated genes in humans, mice, and yeast have identified broad patterns of dosage sharing among daughter paralogs (Qian et al. 2010; Lan and Pritchard 2016), younger, human-specific duplications have yet to be analyzed in this light. Further, no molecular explanations have been provided for the observed expression changes between paralogs.

Great apes have experienced a surge of SDs in the last ∼10 million years, primarily interspersed throughout the genome and potentially contributing to phenotypic differences observed between these closely related species (Prado-Martinez et al. 2013). Human-specific SDs (HSDs), which arose in the last ∼6 million years following the split of the human and chimpanzee lineages, contain genes that have compelling associations with neurodevelopmental features (Charrier et al. 2012; Dennis et al. 2012; Florio et al. 2015; Fiddes et al. 2018; Suzuki et al. 2018; Heide et al. 2020) and disorders (Dennis and Eichler 2016; Dennis et al. 2017; Ishiura et al. 2019). Historically, such young duplications have been poorly resolved in genome assemblies due to their high sequence similarity. Recent sequencing efforts targeted to HSDs have generated high-quality assemblies for many of these loci (Steinberg et al. 2012; Antonacci et al. 2014; O’Bleness et al. 2014; Dennis et al. 2017) resulting in the discovery of at least 30 gene families containing >80 paralogs uniquely duplicated in humans. Most derived HSD genes encode putatively functional proteins and exhibit divergent expression patterns relative to ancestral paralogs across numerous primary tissues, despite HSDs being nearly identical by sequence (on average ∼99.5%) (Dennis et al. 2017). Although there are examples of HSD genes exapting novel promoters and exons at the site of insertion (Dougherty et al. 2017), this cannot explain expression divergence that exists among whole-gene duplications. Differential regulation may be intertwined with associations of species-specific active chromatin modifications at SD loci (Giannuzzi et al. 2014) but historical reference errors and computational challenges in short-read mapping to highly-similar sequences has resulted in poorly annotated epigenetic information at duplicated loci (Chung et al. 2011; Ebbert et al. 2019).

In this study, we characterized patterns of regulatory divergence observed for HSD genes between humans and chimpanzees by quantifying cross-tissue conservation of orthologous gene expression. We found that even the youngest of duplicate genes have diverged in expression and, by comparing expression divergence between ancestral and derived paralogs, have begun to infer functional HSD genes. We leveraged genomic and epigenomic data from hundreds of human lymphoblastoid cell lines (LCLs) to identify differentially expressed (DE) ancestral-derived gene pairs and examined potential molecular contributors to paralogous expression divergence, including copy-number (CN) variation, post-transcriptional regulation, and *cis*-regulatory changes. Finally, we surveyed the active chromatin “landscape” for HSDs by reanalyzing ENCODE histone-modification chromatin immunoprecipitation sequence (ChIP-seq) data, produced a novel “longer-read” ChIP-seq dataset to improve the unique alignment rate in SDs, and functionally validated candidate *cis*-regulatory elements (cCREs) via a reporter assay. Overall, our work demonstrates that *cis*-regulatory divergence, among other mechanisms, drives differential expression following gene duplication and that useful regulatory information can be rescued from existing datasets for duplicated loci.

## RESULTS

### Conservation of HSD gene expression following duplication

To assess the evolutionary trajectory of recent human duplicated genes, we quantified expression of 75 HSD genes from 30 gene families for which high-confidence sequences were available (Dennis et al. 2017) (Table S1). Unlike previous work examining tandem gene duplicates in humans (Lan and Pritchard 2016), the SDs comprising these genes duplicated in an interspersed manner on the same chromosomes, often hundreds of kilobases away, with only two of the 30 gene families residing on separate chromosomes. Each HSD gene family corresponded to a single-copy chimpanzee ortholog and multiple (2–4) human paralogs. If known, we classified the human paralog syntenic with the chimpanzee gene as ancestral and the human-specific paralog(s) as derived (Figure 1A, Table S1). To interpret the evolutionary fate of these genes, we compared expression of HSD paralogs (individual or summed) to chimpanzee orthologs using mRNA-sequencing (RNA-seq) data from three cell lines and four primary tissues (Khan et al. 2013; Pavlovic et al. 2018; Marchetto et al. 2019; Blake et al. 2020) using a lightweight mapping approach that shows high accuracy for paralogous genes (Soneson et al. 2015; Patro et al. 2017). Derived HSD paralogs tended to exhibit lower expression than the chimpanzee ortholog, summed family expression was mostly higher, and ancestral paralogs were less likely to be DE (9/21 expressed ancestral genes showed no differential expression across all cell/tissue types versus 6/37 of expressed derived genes; *p*=0.028, Fisher’s Exact Test) (Figure 1B, Table S2). This suggests that ancestral genes most likely retain their functions, while derived paralogs diverge or pseudogenize.

**Figure 1.**
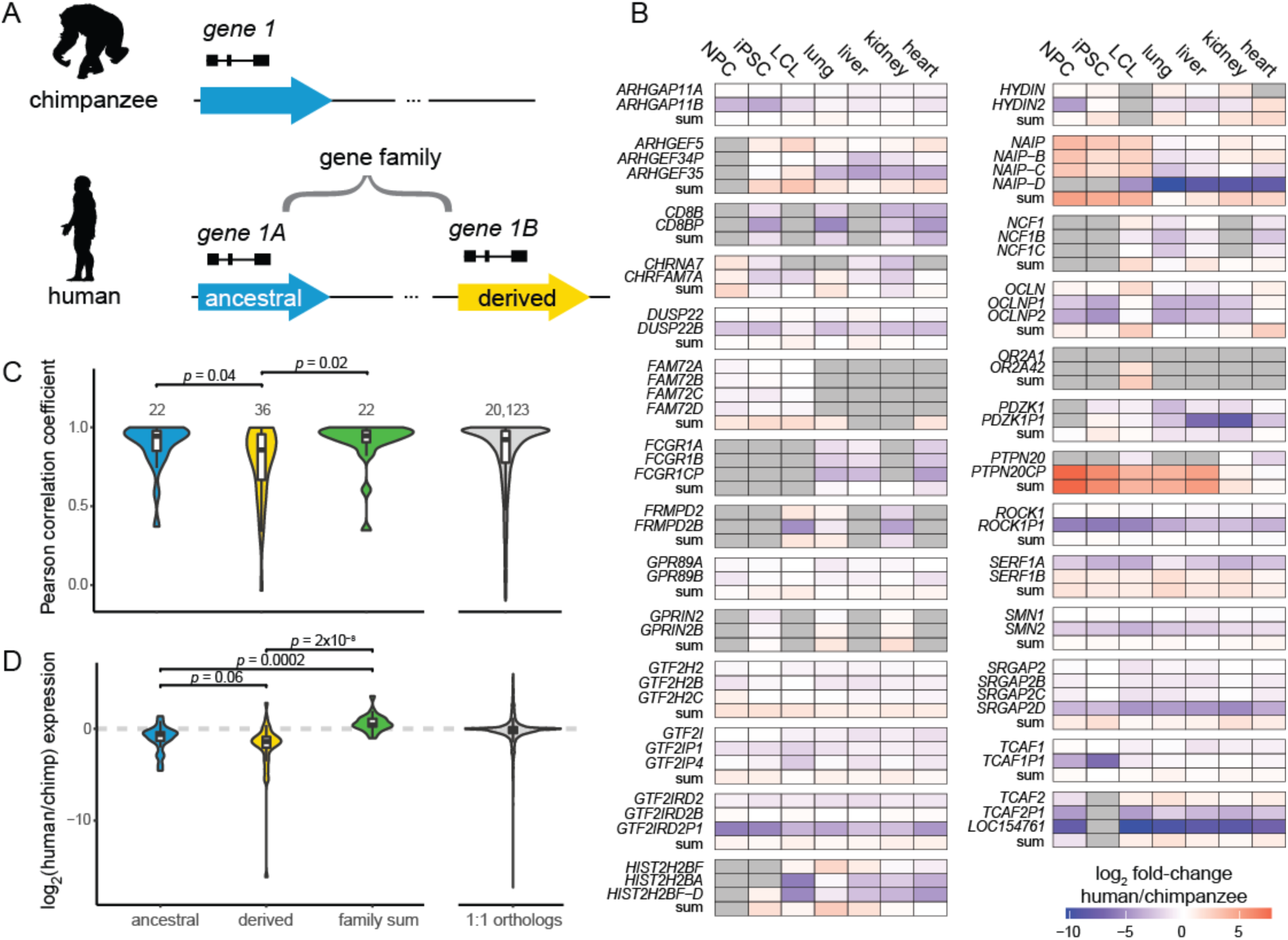
Expression patterns of HSD genes between species. **(A)** Illustration of genes residing within HSDs; the ancestral paralog (blue) corresponds to the chimpanzee ortholog, while derived paralogs (yellow) are human-specific. The ancestral and derived genes comprise a gene family. **(B)** HSD gene expression differences between humans and chimpanzees in three cell lines and four primary tissues. Cells are colored by the log_**2**_**-**fold change of human versus chimpanzee expression. Gray cells indicate non-expressed genes. Differential expression results are provided in Table S4. **(C, D)** Comparison of human gene expression with chimpanzee orthologs. Violin and box plots represent cross-tissue expression correlations (C) and relative expression levels (log_2_ ratio of human versus chimpanzee expression) (D). HSD genes of known evolutionary status were classified as ancestral (blue) or derived (yellow) and compared with the aggregated gene family expression (green). *P*-values were calculated from Dunn’s post-hoc test of a Kruskal-Wallis test. Expression correlations of one-to-one orthologs are visualized for reference.

We next considered expression correlation across the four tissue types and three cell lines as a proxy for expression conservation between human genes and their chimpanzee orthologs. Our expectation was that in the case of subfunctionalization, the summed expression of all HSD paralogs would correlate best with chimpanzee expression, while all individual paralogs would be less correlated; and in the cases of pseudogenization or neofunctionalization, a single paralog would exhibit high correlation with chimpanzee expression (Braasch et al. 2018; Sandve et al. 2018). We found that derived HSD paralogs exhibited significantly lower expression conservation than ancestral paralogs or summed expression, which were statistically equivalent (Kruskal-Wallis test followed by Dunn’s test, Benjamini-Hochberg adjusted *p*<0.05; Figure 1C). This pattern is broadly consistent with maintenance of the ancestral paralog and divergence of expression patterns of the others via pseudogenization or neofunctionalization. Further, the most conserved gene in each family was usually the ancestral paralog (14/22 of known status, *p*<0.001, hypergeometric test). Nevertheless, eight derived paralogs did show strongest conservation of expression with chimpanzee orthologs, including *HIST2H2BA, OR2A42*, and *SERF1B*, and represent candidates of new duplicates usurping functions of their ancestral gene, or the ancestral gene gaining novel expression patterns. We also considered relative expression levels between species and found that across tissues, ancestral paralogs trended toward higher expression than derived paralogs (Kruskal-Wallis test followed by Dunn’s test, Benjamini-Hochberg adjusted *p*=0.058; Figure 1D). As expected, summed HSD expression was significantly higher than ancestral or derived paralogs alone. Taken together, these results indicate uneven maintenance of expression by ancestral HSD paralogs, which is not consistent with subfunctionalization via dosage sharing among daughter paralogs for most gene families.

These results are concordant with our previous finding that derived paralogs globally show a reduction of expression relative to ancestral paralogs, with some exceptions, across diverse human tissues and cell lines from the Genotype-Tissue Expression project (Dennis et al. 2017). To validate this with a strict alignment-based approach, we used long-read PacBio isoform sequencing (Iso-Seq) data, which maps to paralogous loci with higher confidence, from a panel of 24 human biosamples and cell lines (Encyclopedia of DNA Elements (ENCODE) project). From this, we again found globally reduced expression of derived paralogs: 21/41 derived genes were expressed at a level below their ancestral paralog, while two derived genes were higher (p<0.05, Wilcoxon Signed-Rank test with Benjamini-Hochberg correction; Figure S1). Though results should be interpreted cautiously given the low read depth and small number replicates for each biosample, we also observed certain derived paralogs exhibit greater expression than the ancestral paralog in individual tissues or cell types; one compelling example was diverged expression of *ARHGAP11B* in excitatory neurons, which matches published findings related to the novel function of this gene in the human cortex (Florio et al. 2015; Kalebic et al. 2018; Heide et al. 2020).

### Expression of HSD paralogs in lymphoblastoid cell lines (LCLs)

We next focused on LCLs to gain a more detailed understanding of HSD expression patterns across hundreds of individuals with matched genomic data. We estimated transcript abundance using RNA-seq data from 462 human LCLs (Lappalainen et al. 2013) (Table S3) and found high concordance with expression estimates from Iso-Seq data from the LCL GM12878 (Pearson’s *r*=0.94 for 72 genes common to both analyses). We determined that over half (43/75) of HSD paralogs were expressed above one transcript per million (TPM), with the most highly expressed genes including *ARHGAP11A; ROCK1;* the adjacent *GTF2I* and *NCF1* families; and the *DUSP22* family, whose derived paralog *DUSP22B* is missing from the human reference (GRCh38) (Dennis et al. 2017). Comparing expression profiles within gene families, derived and ancestral paralogs globally showed divergent expression levels. In families with at least one expressed gene, all 31 derived genes showed significant differences from their ancestral counterpart, with a median TPM difference greater than two-fold observed in 20 of these. As was found across other cells/tissues, in most cases (25/31) the derived gene had lower expression, which we confirmed for three highly expressed gene families with RT-qPCR and Iso-Seq data (Figures 2A, S2, and Table S4). We noted that some paralogs exhibited clusters of outlier values for derived/ancestral expression ratios that could not be reconciled as copy number (CN) or population of origin differences (Figure S3). Altogether, these results indicate that paralogous HSD genes show divergent expression patterns in LCLs.

**Figure 2.**
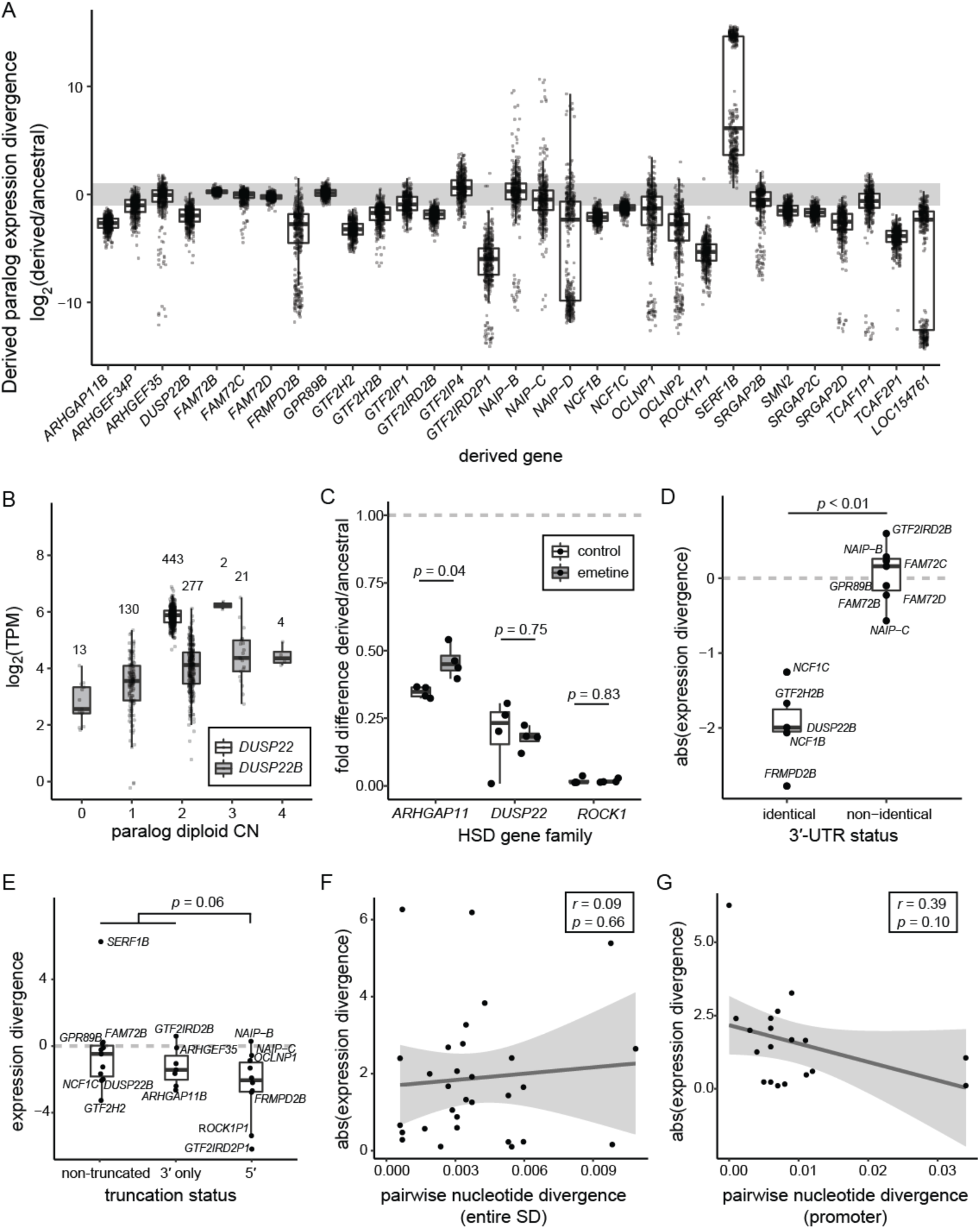
Differential expression of HSD genes in human LCLs. **(A)** Expression divergence of derived genes is plotted as the log_2_ ratio of median derived and ancestral expression for families with at least one LCL-expressed paralog. Each point represents an LCL from the Geuvadis consortium (total N=445) (Lappalainen et al. 2013). The gray bar indicates a two-fold expression difference. **(B)** Expression values of ancestral *DUSP22* (white) and derived *DUSP22B* (gray), stratified by CN. The number of individuals represented in each CN category is denoted over each boxplot. **(C)** Derived/ancestral fold-differences in expression determined from paralog-specific qPCR in control (white) and NMD-inhibited (gray) LCLs (N=4). Statistical significance in panels **C–E** was assessed with a Wilcoxon signed-rank test. **(D)** Absolute value of expression divergence of ancestral-derived gene pairs, stratified by identical or non-identical 3′ UTRs. (E) Comparison of expression divergence across truncation status for all expressed ancestral-derived gene pairs. **(F, G)** Scatterplot of the absolute value of expression divergence versus pairwise nucleotide identity for all expressed ancestral-derived gene pairs for whole duplicons **(F)** and promoters **(G)**. Regression lines (black) and 95% confidence intervals are shown, along with the Pearson correlation coefficient (r) and significance of the regression slope (*p*).

### CN variation and HSD expression

While the genes in this study were chosen for being nearly fixed in modern human populations (Dennis et al. 2017), SD loci are known to be subject to recurrent rearrangement and consequently exhibit varying degrees of CN polymorphism. Understanding that CN variation can alter gene expression levels (Stranger et al. 2007), we sought to characterize its impact on differential expression of HSD genes. After performing paralog-specific CN genotyping (Shen and Kidd 2020) of a subset of individuals for which 1000 Genomes Illumina sequences were available (N=445), we found gene expression was positively associated with CN in about half (28/55) of genes in expressed families (Table S5), indicating that higher CN often but not always translates to increased expression. Notably, derived genes tended to have higher CN (1.3-fold on average, across all assayed genes and individuals) but lower expression overall. We next used linear regression to remove the effect of CN from these comparisons and found 23/25 derived paralogs were still DE with respect to the ancestral (six were not tested due to paralog-specific effects of CN; Table S4). For example, while expression of *DUSP22B* was significantly associated with CN, these effects were insufficient to explain DE relative to *DUSP22* (Figure 2B). Thus, while CN differences alter the mRNA abundance of HSD paralogs, they do not provide an explanation for overall DE of these genes.

### Post-transcriptional regulation of HSD genes

In order to determine if paralogous expression differences are driven by post-transcriptional regulation, we next considered whether HSD transcripts were being processed as nonfunctional pseudogenes. In this scenario, paralogs might be equally transcribed but differentially subject to degradation via nonsense-mediated decay (NMD). To test this, we compared gene expression using available RNA-seq data from human NMD-deficient LCLs (N=4) against controls (N=2) (Nguyen et al. 2012) and found no HSD genes among identified DE genes. We also assessed directly if the ratio of derived to ancestral expression changed for each HSD gene family between NMD-deficient LCLs and controls and found no significant differences, though sample sizes were likely limiting (Figure S4). This result was largely recapitulated by paralog-specific RT-qPCR for three DE HSD genes families (*ARHGAP11, DUSP22*, and *ROCK1*) in four LCLs treated with the NMD-inhibiting drug emetine. Ratios of *ROCK1P1/ROCK1* and *DUSP22B/DUSP22* expression were unaltered by emetine treatment, while *ARHGAP11B*/*ARHGAP11A* expression ratio increased closer to one, consistent with NMD affecting *ARHGAP11B*, though not completely ‘rescuing’ derived expression levels to equal that of the ancestral (Figure 2C). *ARHGAP11B* is a 3′ truncation of *ARHGAP11A*, potentially explaining differences in transcript stability. Altogether, these results suggest that while NMD may alter steady-state expression levels of some HSD genes, it is not a primary driver of their differential expression.

We also examined HSD 3′ untranslated regions (UTRs) for recognition sites of miRNAs expressed in LCLs (Lappalainen et al. 2013) (N=13 3′ UTRs of expressed gene families; mean 94 binding sites per UTR) using TargetScan (Agarwal et al. 2015). Although miRNA binding sites were nearly identical between paralogs, we unexpectedly observed significantly greater expression divergence between paralogs with identical 3′ UTRs (N=5) from those that differed (N=7) (Wilcoxon signed-rank test p<0.01, Figure 2D). While these data cannot rule out a role for miRNAs in HSD transcriptional regulation, this mechanism does not explain observed differential expression of expressed gene families with identical 3′ UTRs, such as *DUSP22* and *NCF1*.

### Role of *cis*--regulation in HSD differential expression

We next aimed to determine if *cis*-regulatory changes contribute to expression divergence of HSDs. Because SDs often generate gene truncations and fusions with adjacent transcribed sequences (Dougherty et al. 2017), we reasoned that gains or losses of promoters or UTRs would likely cause large changes in gene expression. We compared relative expression by truncation status (5′-, 3′-, or non-truncated) of all derived genes in expressed families to their ancestral paralogs. Ancestral and derived genes had more similar expression levels in non-truncating duplications, while truncated genes tended to be less expressed than their ancestral paralogs, particularly 5′ truncations compared to all other HSD genes (*p*=0.057, *t*-test; Figure 2E), in concordance with previous findings (Dougherty et al. 2018). While we may have limited power to detect differences with our small number of genes, these results hint that promoter activity is an important determinant of differential expression patterns. Considering sequence-level changes more broadly, however, we observed no relationship between expression divergence and pairwise nucleotide divergence across entire duplicons or within promoters (Figure 2F–G).

Given that the vast majority of paralog-specific variants (PSVs) distinguishing HSDs are unlikely to be functional, we used publicly available chromatin immunoprecipitation sequencing (ChIP-seq) datasets from the ENCODE project (Consortium and The ENCODE Project Consortium 2012; Davis et al. 2018) to identify likely CREs (H3K4me3, H3K4me1, H3K27ac, and RNA PolII) in a single LCL for which a wealth of functional genomic data exists (GM12878). In each data set, we observed a lower density of bases covered by peaks in SDs (>90% similarity) and HSDs (>98% similarity) compared to randomly sampled regions of equivalent size (empirical p=0.001, N=1000 replicates; Figure 3, in yellow). We posit, as others have previously (Chung et al. 2011; McVicker et al. 2013; Giannuzzi et al. 2014), that this discrepancy is an artifact of the high sequence similarity of SDs, with reads originating from these regions often discarded when mapping to multiple locations of the genome.

**Figure 3.**
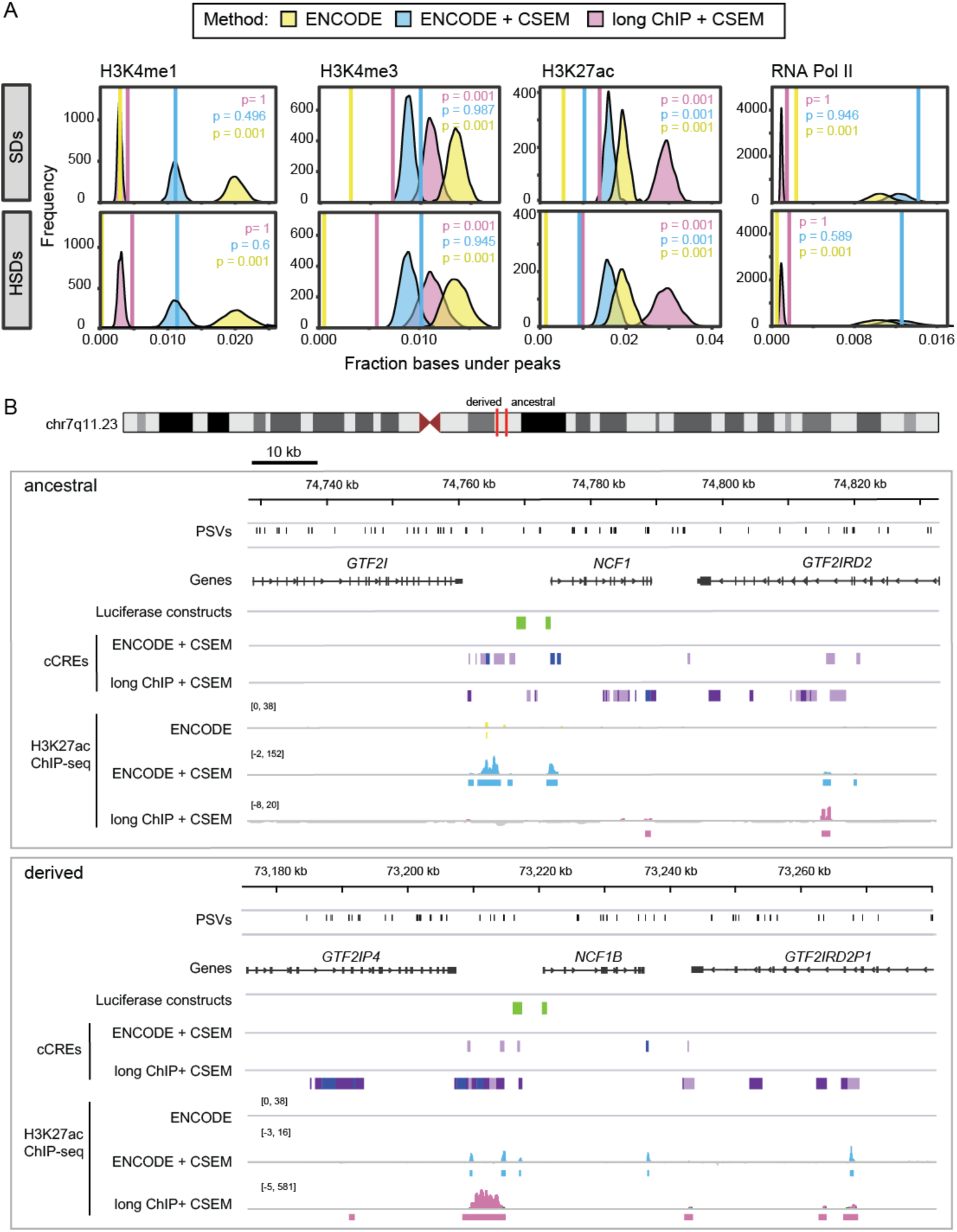
Depletion and recovery of ChIP peaks in SDs. **(A)** The fraction of bases covered by peaks (solid vertical line) was computed in SDs (top) and HSDs (SDs >98% sequence identity, bottom) for three ChIP-seq peak discovery approaches: publicly-available ENCODE peaks (yellow), peaks from multi-mapping and CSEM allocation of ENCODE raw data (blue), and peaks from multi-mapping and CSEM allocation of large-insert ChIP-sequencing (“long ChIP”) data from this publication (magenta). SD coordinates were permuted 1000 times within the human reference (GRCh38), and an expected distribution of the fraction of bases covered was generated. Empirical one-sided *p*-values for depletion are indicated in each graph. **(B)** Chromatin landscape at the chromosome 7q11.23 HSD locus. The ancestral locus (top) and one of its derived loci (bottom) are shown with paralog-specific variants (PSVs) (black), genes (gray), and luciferase-tested regions (green). Candidate *cis*-regulatory elements (cCREs) were identified with an 8-state ChromHMM model of GM12878 H3K4me3, H3K4me1, and H3K27ac data from multi-mapping reanalysis of ENCODE and long ChIP data after CSEM allocation (enhancer states in light and dark purple and promoter states in blue). H3K27ac ChIP-seq data (signal and peak calls) are also shown in yellow, blue, and magenta for published ENCODE, reanalyzed ENCODE + CSEM, and long ChIP, respectively.

To recover this missing information, we implemented a pipeline that allowed reads to align to multiple locations in the genome and then, using CSEM (Chung et al. 2011), iteratively weighted alignments based on the nearby unique mapping rate. Selecting the most likely alignment to allocate a read (i.e., mapping position with the highest posterior probability), we improved peak discovery in SDs and HSDs for the aforementioned chromatin features, erasing the depletion for all but H3K27ac, which was still substantially improved (Figure 3, in blue). The peaks we discovered largely overlapped with the ENCODE peaks, though RNA PolII had a large proportion of peaks unique to our multi-mapping analysis (Figure S5). Using this new dataset, we observed greater enrichment of H3K27ac at the ancestral *DUSP22* versus *DUSP22B*, which we verified at three PSVs using ChIP-qPCR (1.1–2.9-fold difference of ChIP signal; 1.1–2.9-fold difference of dCt values) (Figure S6A). We also noted a correlation of *DUSP22*/*DUSP22B* expression divergence (Pickrell et al. 2010) (Table S3) and differential H2K27ac enrichment at these PSVs (Figure S6B). These findings suggest that reanalysis of ChIP-seq data can accurately identify enriched regions at HSD loci, uncovering potentially divergent regulatory environments.

### Improved peak discovery using longer-read ChIP-seq

To improve our ability to align reads accurately to specific paralogs, we generated longer-read (∼500 bp insert size, 2×250 bp PE Illumina) ChIP-seq (“long ChIP”) libraries (H3K4me3, H3K27ac, H3K4me1, and RNA PolII) from the LCL GM12878. Longer reads mapped to SDs with greater accuracy (Figure S7A), allowing for higher-confidence discovery of novel peaks in duplicated regions using standard single-site mapping approaches. However, all marks except H3K4me1 were still depleted for peaks in SDs relative to the rest of the genome. Subsequently, we analyzed the long ChIP data allowing for multiple alignments and probabilistically assigned reads to one position (Bowtie and CSEM, Figures 3 and S7B). Long ChIP showed increased posterior assignment probabilities with respect to the short-read ENCODE data (Figure S7B), and the depletion of peaks in SDs was erased for H3K4me3, H3K4me1, and PolII (Figure 3, in pink). Notably, for most libraries, fewer overall peaks were identified with long ChIP versus ENCODE data, though the peaks that did exist were largely replicated (on average, 73% of long ChIP peaks corresponded to an ENCODE peak (Chikina and Troyanskaya 2012); Figure S8). Long ChIP peaks tended to be larger with 2.4–3.7 times as many bases per peak except H3K4me1, which had slightly smaller peaks.

To identify putatively functional *cis*-regulatory regions within HSDs, we integrated our reanalyzed ENCODE and long ChIP data into two 8-state chromHMM models (Ernst and Kellis 2012), from which we identified active promoter- and enhancer-like states that we considered to be cCREs (Figure S9). This generated a novel set of cCREs in SDs, as virtually no information is available in the current ENCODE release for these loci (Figure 3B). Because derived gene expression is broadly lower than ancestral, we quantified the proportion of cCREs covering HSDs in 100-kb windows and observed no significant differences between ancestral and derived loci (defined in Dennis, 2017) (Wilcoxon rank-sum test; Figure S10A∗B). We also observed no differences in the fraction of bases covered between ancestral and derived regions in individual ChIP-seq datasets: H3K27ac, H3K4me3, H3K27ac (data not shown), and heterochromatic H3K27me3 domains (Figure S10C). Thus, explanations beyond the overall abundance of chromatin features are needed, as important functional changes in CRE activity may not be reflected in global differences. We did identify differences in presence or absence of individual cCREs, suggesting a more nuanced approach is necessary in pinpointing mechanisms contributing to paralogous expression differences (Figure 3B).

### Impact of non-duplicated regions on HSD gene regulation

HSDs are often transposed many thousands of kilobases from their ancestral loci, and in some cases to different chromosomes. As such, we sought to understand if cCREs outside of our duplicated regions might contribute to paralog-specific regulatory patterns. To do this, we considered physical contacts generated by chromatin looping of HSD promoters with cCREs outside of HSD regions. Using loops identified in GM12878 from promoter capture Hi-C (Mifsud et al. 2015) and H3K27ac HiChIP (Mumbach et al. 2017; Juric et al. 2019), we identified 352 and 26 promoter-interacting regions, respectively (mean size ∼5 kb). We found 59 ENCODE multi-mapping and 106 long ChIP cCREs interacting with an HSD gene promoter. For instance, a chromatin loop connects the *ARHGAP11A* promoter with a cCRE overlapping its non-duplicated 3′-UTR (Figure 4A). The majority (>90%) of promoter-interacting regions reside outside of HSDs, in part due to limitations of Hi-C analysis across duplicated loci (Zheng et al. 2019). These findings indicate that proximal non-duplicated loci may play a regulatory role in regulating duplicated genes.

**Figure 4.**
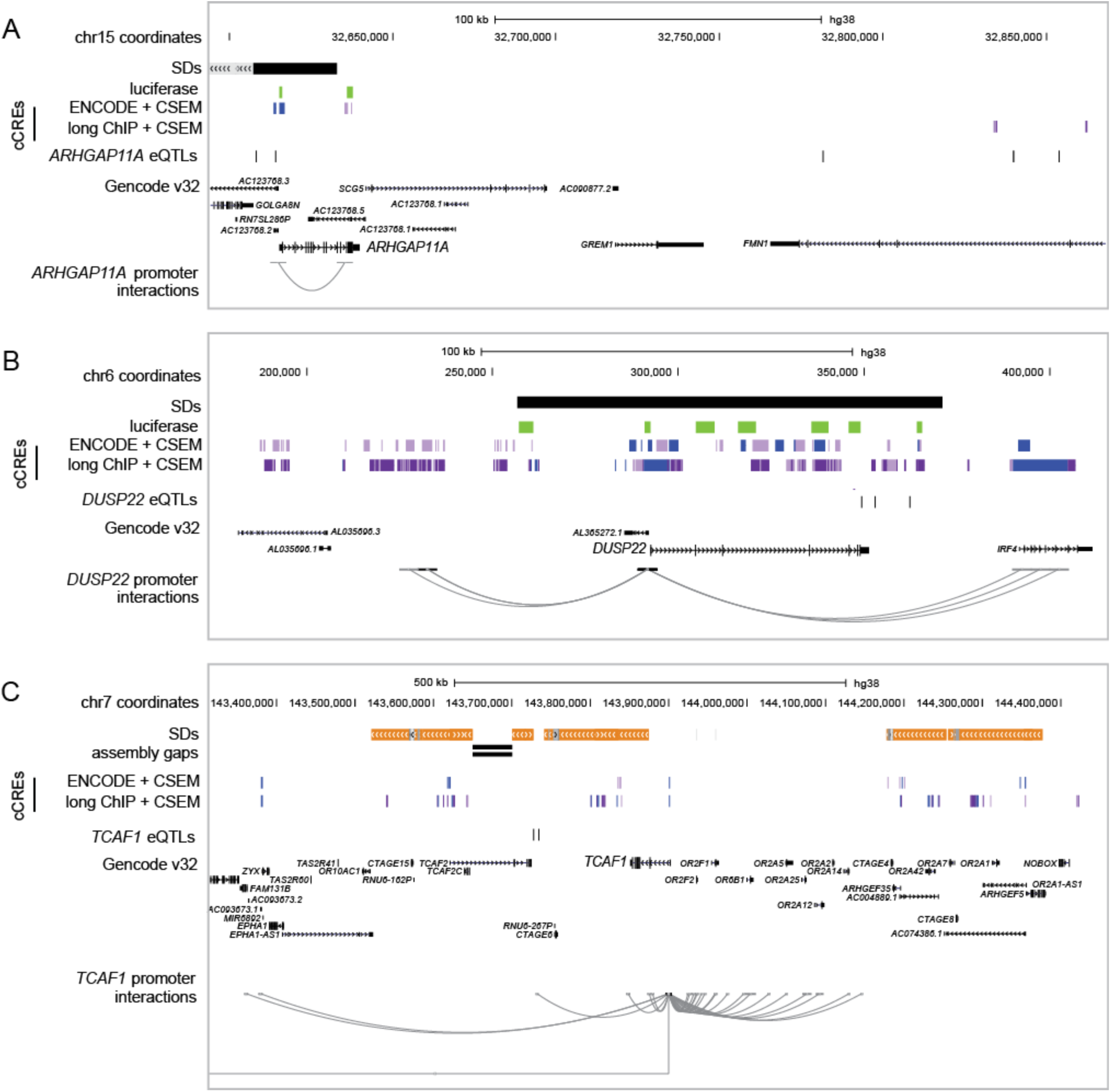
HSD gene regulation in adjacent, non-duplicated sequence. Additional regulatory features were examined in the vicinity of the HSD loci, including **(A)** *ARHGAP11A* at chromosome 15q13.1, **(B)** *DUSP22* at chr6p25.3, and **(C)** *TCAF1* at chr7q35. In each panel, SDs are depicted as gray (>90% identical), orange (>98% identical), and black (unannotated) bars. cCREs as defined in this publication are shown in light and dark purple (active enhancer states 1 and 2) and blue (active transcription start site), with luciferase-tested regions in green. eQTLs defined in this publication and regions previously found to interact with HSD promoters are shown for focal genes. Data were visualized in the UCSC Genome Browser (GRCh38).

We next performed expression quantitative trait locus (eQTL) mapping of HSD genes using our reanalyzed RNA-seq data and existing variant calls from the 1000 Genomes Project (N=460) (Genomes Project, Consortium et al. 2015). From this, we identified 40 HSD genes with significant eQTLs, an increase of 1.5- to 4-fold from published work (Lappalainen et al. 2013; Wen et al. 2015). These eQTLs consisted of 3,279 variants in 8,774 gene-variant pairs. A majority (68%) of eQTLs were located within annotated SDs, but variants identified within SDs are often unreliable (Hartasánchez et al. 2018; Ebbert et al. 2019). Accordingly, we focused on the 1,049 eQTLs in SD-proximal non-duplicated regions and found 439 of them had single-gene associations. For example, four variants were associated with *ARHGAP11A* expression (Figure 4A), while none were identified for *ARHGAP11B* located ∼2 Mb proximal to its ancestral locus. Similarly, four eQTLs were identified for *DUSP22* on chromosome 6 (Figure 4B), though all were located in an SD, while 26 variants were linked with the derived paralog *DUSP22B* on chromosome 16. We intersected SD-proximal eQTLs with our cCREs, reasoning that functional elements would be sensitive to genetic variation and, thus, contain eQTLs. We found that five ENCODE multi-mapping and 15 long ChIP cCREs contained an HSD eQTL. Finally, 169 eQTLs fell within loci showing significant Hi-C interactions with HSD promoters (31 of these regions, total size ∼160 kb). For instance, the *TCAF1* promoter interacts with a region ∼170 kb downstream that is near two SNPs associated with *TCAF1* and *TCAF2* expression (Figure 4C). Altogether, these findings highlight the potential for adjacent, unique sequences to drive divergent regulation of HSDs genes.

### Differential activity of *cis*-acting elements between paralogs

Using our combined datasets, we examined three HSD loci containing gene families expressed most highly in LCLs (*ARHGAP11, NCF1*, and *DUSP22*) to identify functional changes in CREs that may contribute to paralogous expression divergence (Figures 5, S11–S13). In all three cases, the ancestral paralog exhibited significantly greater expression compared to derived paralog(s) (Figure 5A). To determine if sequence differences within CREs identified from our chromHMM annotations were sufficient to drive differences in gene expression, we performed luciferase reporter assays on paralogous promoters and enhancer candidates in HeLa cells and LCLs.

**Figure 5.**
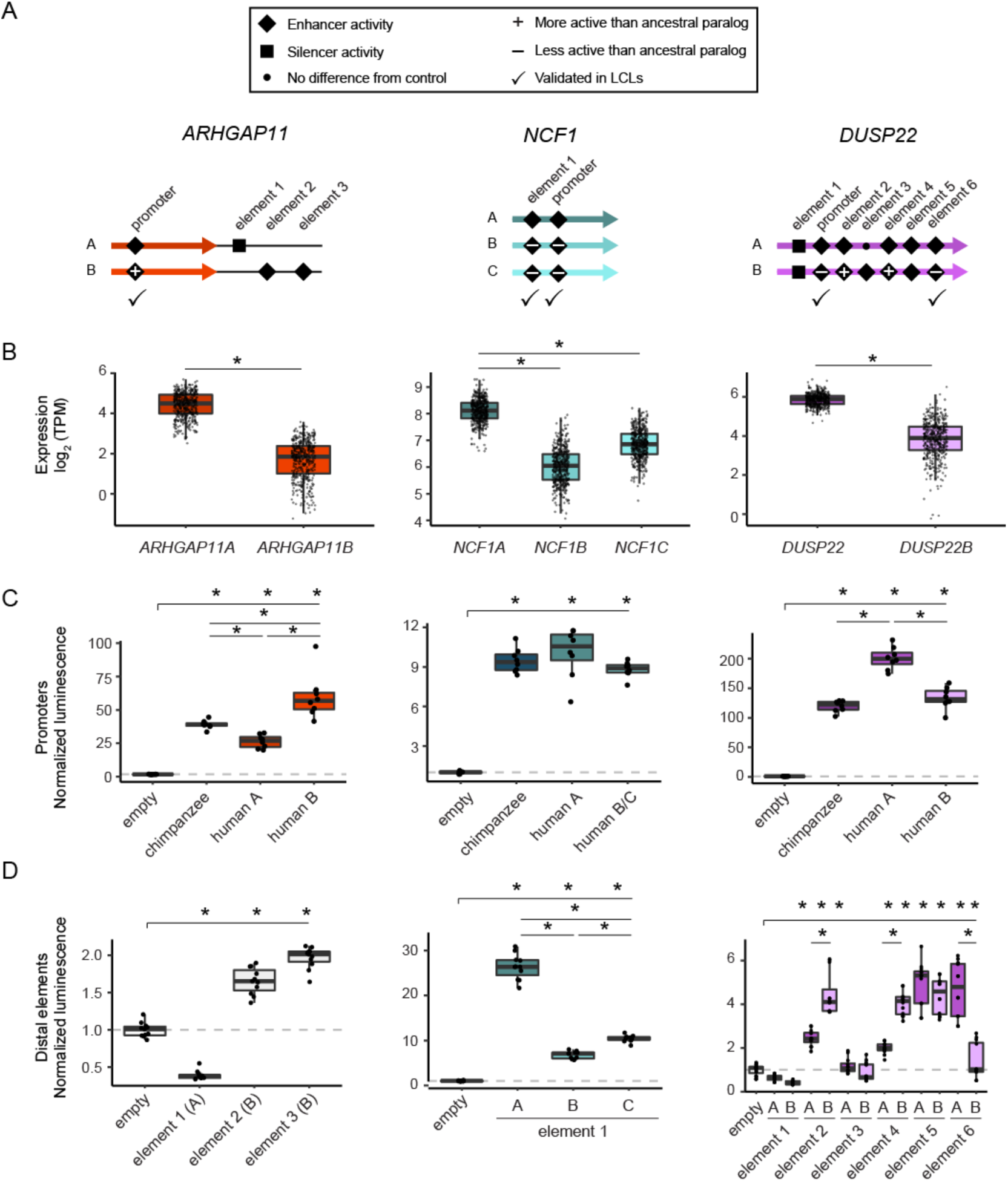
Functional characterization of cCREs in HSDs. cCREs (putative promoters and enhancers) from three HSD duplicate gene families (*ARHGAP11, NCF1*, and *DUSP22*) were tested in luciferase reporter assays for activity. **(A)** Cartoons indicating the relative locations of each candidate sequence within or adjacent to HSDs (thick, colored arrows). All experiments (Figures S14∗S15) are summarized as follows: inactive sequences are shown with a small dot, enhancer sequences are shown with a diamond, and silencer sequences are shown with a square; differentially active derived sequences (relative to ancestral) are marked with a plus or minus sign; elements tested (and validated) in LCLs are indicated with a check mark. **(B)** mRNA levels (TPM) for the three tested HSD gene families in human LCLs (N=445). **(C)** Representative luciferase reporter experiments for promoters of the paralogous HSD genes and orthologous chimpanzee sequences in HeLa cells. Significantly different activity (*p*<0.05, Tukey’s post-hoc test of ANOVA) from the negative control is indicated along the top bar over each panel, and significant differences among homologous sequences are indicated between boxplots. The *p*-values for each comparison for all experiments are available in Table S6. **(D)** Representative luciferase reporter experiments for candidate enhancers from the same gene families in HeLa cells, with significant activity over/under baseline indicated along the top bar, and significant differences between paralogous sequences between boxplots (*p*<0.05, Tukey’s post-hoc test of ANOVA).

#### ARHGAP11A/ARHGAP11B

The promoter of *ARHGAP11B* exhibited greater activity compared to the chimpanzee ortholog and ancestral paralog in both HeLa and LCLs (∼4-fold difference in activity between HSD paralogs, *p*<5×10^−10^ in both cell lines; Figures 5B, S14A, S15A). This was in contrast to mRNA levels in LCLs, where the ancestral *ARHGAP11A* was more highly expressed. With no CREs identified within the shared *ARHGAP11* HSD, we posited that distal elements may drive differential expression between these paralogs. We identified putative enhancers unique to each paralog outside of the shared HSD, which comprised one downstream of the *ARHGAP11A* duplicon (that was also found to interact with the promoter from our Hi-C analysis) and two downstream of *ARHGAP11B*. In HeLa cells, the *ARHGAP11A* element showed weak repressive activity (0.3-fold difference, *p*<2×10^−16^), while the *ARHGAP11B* elements showed modest activity over baseline (∼2-fold difference, *p*<2×10^−14^ each), leaving the primary driver of differential expression for these genes as yet determined (Figures 5C, S14B).

#### NCF1/NCF1B/NCF1C

Promoters of the ancestral *NCF1* and its derived paralogs *NCF1B* and *NCF1C* genes did not exhibit significant differential activities in LCLs and modest differences in HeLa (0.8-fold difference, p<0.001; Figures 4A, S14A, S15A). However, an enhancer element common to all three paralogs showed the greatest activity for the ancestral *NCF1* paralog in both cell types. This was concordant with differential mRNA levels (∼3-fold difference over either derived in LCL; *p*<0.002 for all comparisons) (Figures 5, S15B). Thus, this enhancer, if targeted to *NCF1* and its paralogs, may contribute to differences in their mRNA levels.

#### DUSP22/DUSP22B

*DUSP22* (ancestral) and *DUSP22B* (derived) promoters showed differential activity concordant with their gene expression in both HeLa and LCLs (i.e., the human ancestral paralog exhibited significantly greater activity than both the human derived and chimpanzee ortholog; ∼1.5-fold difference; *p*<5×10^−13^) (Figures 5B, S14A, S15A). We also tested six putative enhancers shared between the two paralogs in HeLa cells and found four active elements, of which two showed differential activity opposite to that of gene expression and one tracked with differential paralog expression (Figure 5C). We subsequently validated the latter enhancer element in LCLs (∼1.4-fold difference; *p*<2×10^−16^) (Figure S15C). From this, we concluded that the difference in promoter activity is the primary driver of *DUSP22* and *DUSP22B* differential expression, though distal CREs also likely play a role in modulating transcription. Altogether, results from our reporter assays demonstrated that small sequence differences in HSDs can alter *cis*-regulatory activity.

## DISCUSSION

In this work, we provide evidence that recently duplicated, human-specific genes exhibit differential expression at least in part due to divergent *cis*-acting regulation. Historically, these regions have been poorly characterized genetically and epigenetically. Here, we performed an in-depth analysis of gene expression using tissues and cell lines from humans and chimpanzees and delved into putative regulatory mechanisms contributing to paralogous expression divergence, including contributions of CN variation, post-transcriptional modifications, and alterations of CREs via re-analysis of histone ChIP-seq data.

Although previous work (Dennis et al. 2017; Dougherty et al. 2017) quantified HSD gene expression across human cells/tissues, we expanded our analysis to consider all annotated isoforms for each HSD gene, improved or created annotations for derived HSD genes, and orthogonally validated results using Iso-Seq data. By comparing expression of human and chimpanzee homologs, we assayed general mechanisms of duplicate gene fates at relatively short evolutionary time scales (<6 million years). In agreement with previous analysis of whole-genome duplications in teleost fishes (Sandve et al. 2018), we found that the dominant scenario for HSD genes involved a single paralog maintaining ancestral gene expression, perhaps due to purifying selection. This is largely consistent with nonfunctionalization or neofunctionalization of the other paralog(s). Some gene families (such as *CD8B, GTF2IRD2*, and *NAIP*) displayed patterns consistent with subfunctionalization, as their summed expression correlated better with that of chimpanzee than any individual paralog; however, in these cases the difference was small (difference in Pearson’s *r* <0.05 between sum and most correlated paralog). Unlike previous work that characterized duplication events hundreds of millions of years old (Braasch et al. 2016; Lien et al. 2016), HSDs arose very recently, suggesting that changes in expression were rapidly acquired. Our lack of evidence for dosage sharing stands in contrast to that of Qian et al. (2010), who reported an inverse relationship between gene expression and number of paralogs in duplicates arising since the split of the human and mouse lineages, as well as the ancient split of the fission and budding yeasts (>300 million years). Additionally, a study by Lan and Pritchard (2016) found that mammalian tandem duplicated genes tend to be co-regulated and subfunctionalized by dosage, although they excluded many of the genes in our study due to high sequence identity; further, most HSD genes are interspersed at least many hundreds of kilobases from each other throughout the human genome. They and others (Rodin and Riggs 2003) suggest that divergence of gene regulation is more likely in the case of large genomic rearrangements, which can immediately place daughter paralogs in novel regulatory environments, such as topological domains, heterochromatin, or transcriptional hubs. We note that even very recent (<1 million years) duplications (Dennis et al. 2017), such as the gene families *DUSP22, SERF1, SMN, TCAF1*, and *TCAF2*, exhibited differential expression between paralogs, suggesting regulatory differences arose rapidly or existed at the time of duplication.

We functionally investigated three of the highest LCL-expressed HSD gene families to better understand how altered CREs may contribute to paralogous expression divergence. Although LCLs may not represent the most biologically relevant system to characterize species-specific features commonly associated with humans, such as anatomical distinctions in brain and musculoskeletal features, there is evidence of immune-related differences across great apes (Barreiro et al. 2010). Further, it has been suggested that humans are more prone to autoimmune disease than chimpanzees, particularly as a result of T and B cell response to viral infection (Anon 2013; Anon 2017). Interestingly, ancestral *NCF1*, which encodes Neutrophil Cytosolic Factor 1, has been shown to regulate autoimmune response via T cell activation in mice (Hultqvist et al. 2004). CN differences of the gene and its derived full-length paralogs (*NCF1B* and *NCF1C*) are associated with reduced risk of systemic lupus erythematosus (Zhao et al. 2017), suggesting an additive functional effect. *DUSP22*, which encodes a tyrosine phosphatase, is also a regulator of immune response, with knockout mice exhibiting increased inflammation and autoimmune encephalomyelitis as a result of enhanced T cell proliferation (Li et al. 2014). The full-length paralog *DUSP22B* is located at chromosome 16p12.1 at variable CN (Figure 2B) but functionally uncharacterized, as it is missing from the human reference. Thus, any information related to gene regulation may provide insights into the putative roles of these HSD paralogs in immune response and related disorders. For example, it is plausible that *DUSP22B* is functionally redundant with *DUSP22* and expressed at variable dosage in humans, in a manner similar to *NCF1*.

Our characterization of promoters and distal regulatory elements of these gene families uncovered potential drivers of differential expression. The ancestral *DUSP22* promoter and an enhancer element upstream of *NCF1* both exhibited activity patterns concordant with the mRNA levels of these gene families for which the ancestral gene is most highly expressed. Searching for potential *trans* effectors, we identified an ancestral *DUSP22* deletion of four bases in a homopolymer repeat, with as many as 13 similar transcription-factor binding sites (TFBSs) found only in the less active *DUSP22B* and chimpanzee *DUSP22* ortholog. Some of these belonged to transcriptional repressors (ZNF394 and ZNF350). Examining predicted TFBSs within *NCF1* promoters, which did not exhibit differential activity, we observed no gains or losses of any TFBSs relative to chimpanzees. Puzzlingly, no predicted TFBSs were unique to the most active *NCF1* enhancer, but the paralogous *NCF1B* and *NCF1C* possessed many TFBSs that were missing from the ancestral, at least one of which belonged to the transcriptional repressor ZNF394. Finally, we showed that the derived *ARHGAP11B* promoter exhibited greater strength in a reporter assay versus the ancestral *ARHGAP11A* ancestral paralog and chimpanzee ortholog in a luciferase reporter assay. We noted a single PSV that more than doubles the number of significant motif matches of the more active *ARHGAP11B* promoter for the transcriptional activators FLI1, GABPA, ETS1, and ELK1 (Figure S16). Based on chimpanzee homology, these are likely *ARHGAP11A*-specific losses, which matches its reduced activity relative to the derived paralog and chimpanzee ortholog (Figure 5). While our reporter-assay results were discordant with the mRNA-expression levels of these paralogs in LCLs, they may help to explain the unique expression of *ARHGAP11B* in other cell types, such as cortical progenitor neurons (basal radial glia) (Florio et al. 2015). This highlights a need to expand our characterization of CREs to additional relevant cell types and systems, such as neurons and/or cortical organoids. Nevertheless, this study is the first to directly validate and compare CRE activities of recently duplicated paralogs in humans pointing to a variety of mechanisms contributing to differential expression in these gene families.

Though this work represents an important step toward a more complete picture of the regulation of HSD genes, there are still some technical limitations to overcome. We implemented an alignment-free approach for transcript quantification, which accommodates ambiguous mappings resulting from methods and is demonstrated to accurately distinguish highly similar transcripts (Soneson et al. 2015; Patro et al. 2017). However, reads originating from multiple transcripts are assigned probabilistically, and, as such, some genes may appear artificially similar in expression. For instance, *DUSP22B* expression was nonzero in individuals completely missing this paralog (Figure 2B) due to the recent nature of this duplication (<1 million years), which has resulted in very few PSVs differentiating the duplicates at the mRNA level. Conversely, RNA-seq quantification of older gene families (such *HIST2H2BF)* appeared entirely distinguishable between paralogs. In addition, the available contigs of complete HSD loci have only been generated from a single haplotype (Dennis et al. 2017). As such, PSVs are not necessarily fixed, and may also be shared between genes (Dumont 2015). Accordingly, a more advanced understanding of genetic variation at HSD loci is required to separate similarly expressed genes with maximum leverage. This would also allow for the identification of reliable eQTLs within duplications. We also find, as expected, that longer read lengths allow greater bioinformatic distinction between duplicated loci, which remains an inherent limitation of many existing data sources, such as the relatively short reads (∼30 nt) of ENCODE ChIP-seq. While longer reads are fundamentally more informative, some discrepancies between the ENCODE and long ChIP CSEM analyses could be a result of misallocation of reads. We were encouraged to find that a standard single-mapping approach (bwa mem) identified novel peaks in HSD, albeit at a lower rate than genome-wide. As such, longer ChIP read lengths can be used in future studies to further describe the epigenetic state of duplicated loci with a broader array of epitopes and cell/tissue types. Finally, though we used existing chromatin conformation information to connect non-duplicated adjacent regions with HSD gene promoters, improved experimental and bioinformatic methods are required to accurately link distal CREs within SDs to their target promoters. Like other genomic assays, available chromatin interactions from Hi-C data are sparse in SD regions due to poor mapping quality at similar paralogs. To our knowledge, one method exists (mHiC) that allocates multi-mapping reads to their possible alignments (Zheng et al. 2019), similar to CSEM. Future work might discover interactions within duplicated regions by reanalyzing chromatin conformation data in the most recent human reference (GRCh38) with this tool.

Despite limitations, we show that HSD regions contain differentially regulated genes and have identified potentially active chromatin in these regions. Using these data, we demonstrated that recently duplicated regulatory sequences are often functional, and that small sequence changes can significantly alter their activity. Our study did not identify a universal factor that gives rise to the observed differential expression between paralogs, and the underlying molecular mechanisms are likely unique to each HSD gene. In the three duplicate gene families tested, promoter activity is only sometimes concordant with overall gene expression, suggesting that other types of regulatory elements, like enhancers and silencers, may cooperatively control overall expression. Currently, the challenge is to pinpoint functional CREs impacted by PSVs or residing within non-duplicated regions that may differentially alter specific paralogs. We have produced and leveraged a variety of analyses to narrow down likely candidates by chromatin state, expression modulation, and physical proximity to promoters. However, the number of candidate regions is too great to test via low-throughput methods such as luciferase reporter assays. This problem is exacerbated by the need to compare regulatory behavior across multiple cell types. To address this, massively-parallel reporter assays should be employed to validate and quantify CRE activity of thousands of candidate paralogous sequences. Such data could determine to what extent HSD gene expression is predicted by nearby CRE activity. We could also integrate additional types of data, such as targeted chromatin capture of CREs within SDs (such as 4C or capture Hi-C) or nascent transcription (GROseq, 5’ CAGE). Finally, characterization of DNA methylation, which is especially challenging in duplicated loci, will be vital to build a more complete picture of the epigenetic landscape of HSD loci. This study represents a first step toward improving quantification of gene expression and active chromatin states in recent duplications and provides a foundation for future work characterizing regulatory and functional changes in recently duplicated loci.

## MATERIALS AND METHODS

### Quantification of HSD gene expression

The following aligned Iso-Seq filtered alignments were obtained from the ENCODE portal (Davis et al. 2018) (https://www.encodeproject.org/): ENCFF225CCJ, ENCFF648NAR, ENCFF192PJS, ENCFF538BNH, ENCFF596ODX, ENCFF694CBG, ENCFF049QGQ, ENCFF846YHI, ENCFF600MGT, ENCFF810FRP, ENCFF479SQR, ENCFF504GVG, ENCFF731THW, ENCFF936VUF, ENCFF939EUU, ENCFF100RGC, ENCFF927MKK, ENCFF292UIE, ENCFF738RAA, ENCFF437SYY, ENCFF989FKA, ENCFF911RNV, ENCFF305AFY, ENCFF016SHE, ENCFF479EHE, ENCFF914XOH, ENCFF054YYA, ENCFF470UHX, ENCFF158KCA, ENCFF757LOZ, ENCFF809QBD, ENCFF779VVX, ENCFF971JDY, ENCFF319JFG, ENCFF901XCR, ENCFF117DUA, ENCFF772MSZ, ENCFF644PGG, ENCFF955XSL, ENCFF509GHY, ENCFF803KIA, ENCFF058HQU, ENCFF973OML, ENCFF745HHL, ENCFF472TSL. Reads were counted per HSD gene with HTSeq (Anders et al. 2015) before calculating RPKM values. For Figure S2B, *DUSP22* and *DUSP22B* reads were counted separately based on PSV sequence using Samtools mpileup and the raw alignments in the following accessions: ENCFF132HLS, ENCFF234YIJ, ENCFF407TMX, ENCFF592BQN, ENCFF615XZM, ENCFF810UWA.

Human and chimpanzee RNA-seq data were quantified alignment-free with a custom reference transcriptome. Due to poor annotation of many HSD paralogs, custom transcriptomes were generated to ensure equivalent isoform models for paralogous genes, biasing against differential expression. First, transcript sequences for ancestral genes were extracted from GENCODE and mapped to derived human loci (contig from (Dennis et al. 2017) or GRCh38 for *SERF1B*) and a long-read chimpanzee assembly (Kronenberg et al. 2018) using BLAT (Kent 2002). GENCODE v27 transcripts were used for human-chimpanzee comparisons, since the chimpanzee transcriptome (Kronenberg et al. 2018) was built on this version; for human-only analyses, GENCODE v32 was used. Alignments were manually curated, and new derived transcripts were extracted from contigs. These transcripts, in addition to HSD transcripts generated from whole-isoform sequencing of brain tissue (Dougherty et al. 2018), were added to GENCODE (human) or the chimpanzee (after aligning to the chimpanzee assembly) transcriptome. Expression quantification was performed using Salmon v1.2.0 (Patro et al. 2017), the custom transcriptomes, and reference genomes (GRCh38 or Kronenberg et al.) as a decoy sequence. For paired-end data, we used the flags “--validateMappings” and “--gcBias”. RNA-seq data were first lightly trimmed prior to quantification using trim_galore (https://github.com/FelixKrueger/TrimGalore) with the following flags: -q 20 --illumina --phred33 --length 20. Length-normalized TPM values or counts per gene were obtained using the tximport package in R (Soneson et al. 2015).

### Differential expression analysis

Human and chimpanzee RNA-seq data from four primary tissues (Blake et al. 2020), LCLs (Khan et al. 2013; Blake et al. 2020), induced pluripotent stem cells (iPSCs) (Pavlovic et al. 2018), and iPSC-derived neural progenitor cells (Marchetto et al. 2019) were analyzed as described above. Count data from chimpanzee genes were duplicated to allow for pairwise comparison to each HSD duplicate, as well as the sum of all HSD genes in each family. Genes expressed below the 75% percentile (corresponding to 1-2 counts per million reads) were filtered from the analysis, leaving 16,752–18,225 genes. A linear model including species and sex was fitted to each shared gene (N=55,461) using limma-voom (Law et al. 2014; Ritchie et al. 2015), and differentially expressed genes were identified at a 5% false discovery rate (FDR) (Nguyen et al. 2012).

### Copy number-controlled differential expression analysis

Paralog-specific CN estimates were generated using QuicK-mer2 (Shen and Kidd 2020), whole-genome sequence data from the 1000 Genomes Project (30X) (Fairley et al. 2020), and a custom reference consisting of GRCh38 plus an additional contig representing the *DUSP22B* duplicon (Dennis et al. 2017). Expression analysis was performed using RNA-seq data from LCLs included in the Geuvadis study (Lappalainen et al. 2013) for which CN genotypes were generated (N=445). Ancestral-derived gene pairs were compared with a linear model to identify significant differences in log_2_-transformed TPM values after controlling for continuous CN genotypes. Models were first fit with an interaction coefficient, and if no interaction was detected (*p* > 0.05), models were fit to expression and CN only. Resulting *p*-values were corrected via the Benjamini-Hochberg procedure using the R package qvalue (http://github.com/jdstorey/qvalue) and used to identify differential expression of ancestral-derived gene pairs at a 5% FDR. For visualization purposes (Figure 2B), *DUSP22* CN genotypes were adjusted to known values for GM12878 (as determined by fluorescence *in situ* hybridization in (Dennis et al. 2017)).

### Cell Culture

Human LCLs were obtained from the Coriell Institute. The cells were grown in suspension in RPMI 1640 medium (Genesee Scientific) supplemented with 15% fetal bovine serum, 100 U/mL penicillin, and 100 µg/mL streptomycin and maintained at 37°C with 5% CO2. To test the impact of NMD inhibition, two million cells of each LCL (GM19204, GM18508, GM19193, GM19238, GM12878, and S003659_Chimp1) were grown overnight and subsequently treated with 100 µg/ml of emetine (Sigma) for seven hours (Noensie and Dietz 2001). Parallel cultures were left untreated and grown at standard conditions. HeLa cells were grown in Dulbecco’s Modified Eagle Medium (DMEM), High Glucose, with L-Glutamine (Genesee Scientific) supplemented with 10% fetal bovine serum (Gibco, Life Technologies), penicillin (100 U/mL) and streptomycin (100 µg/mL) (Gibco, Life Technologies) at 37°C with 5% CO_2_.

### RNA extraction and cDNA generation

LCLs were harvested and added to an appropriate volume of TRIzol® solution (Invitrogen™) (1 ml per 10^7^ cells) and stored at -80°C for ∼24 hr before extraction to ensure complete lysis of cells. The next day, 200 µl of chloroform (Fisher Scientific) was added, and the homogenate was shaken vigorously for 20 seconds and incubated at room temperature for 2–3 min. Samples were spun at 10,000×*g* for 18 min at 4°C and the aqueous phase was transferred to a sterile RNase-free tube. An equal volume of 100% RNAse-free ethanol was added, samples were mixed by vortex, and then purified with an RNeasy Mini Kit (Qiagen). Samples were eluted in 30 µl RNase-free water and stored at -80°C. Transcriptor High Fidelity cDNA Synthesis Kit (Roche) was used for cDNA synthesis with OligodT primers. Following reverse transcription, samples were treated with RNase A (Qiagen) at 37°C, and cDNAs were stored at - 20°C.

### Identification of miRNA binding sites

For ancestral paralogs of each HSD gene family, the 3′-UTR was extracted from canonical transcript isoforms using the UCSC Genome Browser (GRCh38) and compared using blastn (Altschul et al. 1990) against existing alignments of homologs previously generated for human, chimpanzee, and rhesus (Dennis et al. 2017). Using TargetScan 7.0 and annotated miRNA sequences and families (release 7.1; Sept 2016) (Agarwal et al. 2015), we identified miRNA targets of individual human paralogs and non-human primate orthologs.

### Correlation of expression skew and sequence divergence

Ancestral-derived paralog expression skew was calculated as the absolute value of log_2_(derived/ancestral), using the median TPM values for each gene and a pseudocount 1×10^−4^ (an order of magnitude below the smallest nonzero value). Sequence divergence as the pairwise identity with the ancestral sequence was taken from (Dennis et al. 2017). Gene families were included if at least one paralog was expressed at a level >1 TPM. For promoters, sequence divergence was tabulated as the sum of all mismatches and alignment gaps within ±500 base pairs of the transcription start site (Gencode v32). These quantities were correlated and the strength of the relationship was determined with a linear regression.

### ChIP assays

ChIP assays were carried out as previously described with minor modifications (O’Geen et al. 2019). GM12878 cells were cross-linked in growth media containing 1% formaldehyde (Fisher Scientific BP531) for 10 min at room temperature and the reaction was stopped with 0.125 M glycine. Cross-linked cells were washed twice in PBS and stored at -80°C. 2×10^6^ cells were used per ChIP assay. Cells from two biological replicates were lysed with ChIP lysis buffer (5 mM PIPES pH8, 85 mM KCl, 1% Igepal) with a protease inhibitor (PI) cocktail (Roche). Nuclei were collected by centrifugation at 2,000 rpm. for 5 min at 4°C and lysed in nuclei lysis buffer (50 mM Tris pH8, 10 mM EDTA, 1% SDS) supplemented with PI cocktail. Chromatin was fragmented in microTUBEs with the E220 (Covaris) using the low cell shearing protocol (Duty cycle 2%, PIP 105, CPB 200, 4 min) and diluted with 5 volumes of RIPA buffer (50 mM Tris pH 7.6, 150 mM NaCl, 1 mM EDTA pH8, 1% Igepal, 0.25% Deoxycholic acid). ChIP enrichment was performed by incubation for 16 h at 4°C with the following antibodies: 2 µg H3K27ac antibody (Active Motif #39133), 4 µg H3K4me1 antibody (Millipore 07-436), 2 µg H3K4me3 antibody (Active Motif #39915), or 2 µg RNA Polymerase II (PolII) antibody clone 8WG16 (Covance MMS- 126R). RNA PolII samples were incubated for an additional hour with 2 µg Rabbit Anti-Mouse IgG (MP Biomedical #55436). Immune complexes were bound to 20 µl magnetic protein A/G beads (ThermoFisher) for 2 hours at 4°C. Beads were washed 2x with RIPA, 3x with ChIP wash buffer (100 mM Tris pH8, 500 mM LiCl, 1% Deoxycholic acid) and once with ChIP wash buffer plus 150 mM NaCl. ChIP samples were eluted in 100 µl ChIP elution buffer (50 mM NaHCO_3_, 1% SDS) and cross-linking reversed with addition of 0.5 M NaCl and heating at 65°C overnight. Samples were treated with 2 µg RNaseA (Qiagen) and DNA was purified using the QIAquick PCR Purification Kit (Qiagen). ChIP enrichments were confirmed by qPCR with 2× SYBR FAST mastermix (KAPA Biosystems) using the CFX384 Real-Time System C1000 Touch Thermo Cycler (BioRad). ACTB primers served as positive control and HER2 primers as negative controls (Table S7). ChIP enrichment was calculated relative to input samples using the dC_t_ method (dC_t_ = C_t_[HER2-ChIP]-C_t_[input]). Each entire ChIP sample was used to prepare Illumina sequencing libraries using the KAPA Hyper Prep Kit (Roche). Adapter-ligated DNA was separated on a 2% E-Gel EX (Invitrogen) and the 500–800 bp fraction was excised and purified using the QIAquick gel extraction Kit (Qiagen). Indexed primers were used to generate dual-indexed libraries and amplified libraries were size selected (500–700 bp) using the PippenHT (Sage Science). Equimolar library amounts were pooled and sequenced on the NovaSeq SP (Illumina).

### Analysis of ChIP-seq data

ChIP-seq peaks obtained with the ENCODE pipeline were directly downloaded from the ENCODE portal (Davis et al. 2018) (https:www.encodeproject.org/) for H3K4me3 (ENCFF228GWY), H3K27ac (ENCFF367KIF), H3K4me1 (ENCFF453PEP), POLR2A (ENCFF455ZLJ), and H3K27me3 (ENCFF153VOQ). For “short” ChIP-seq peak calling using raw ENCODE data, GM12878 ChIP-seq reads were downloaded from the ENCODE portal for RNA Polymerase II (ENCSR000AKA), H3K4me3 (ENCSR000BGD), H3K4me1 (ENCSR000AKF), H3K27ac (ENCSR000AKC), and H3K27me3 (ENCFF000OBB). Illumina adapters and low quality bases (Phred score < 20) were trimmed using Trimmomatic (Bolger et al. 2014) (parameters SLIDINGWINDOW:4:20 MINLEN:20) and aligned to a custom reference genome (GRCh38 with an added *DUSP22B* contig) using single-end Bowtie (Langmead et al. 2009) configured to allow multiple mappings per read (parameters a -v2 -m99). After mapping, PCR duplicates were removed using Picard Markduplicates and secondary alignments were removed with samtools v1.9. Multi-mapping reads were allocated to their most likely position using CSEM v2.4 (Chung et al. 2011). CSEM was run using the --no-extending-reads option and the fragment size was calculated with phantompeakqualtools run_SPP.R script (Landt et al. 2012). A custom script was developed to select the alignment with the highest posterior probability as assigned by CSEM for each multi-mapping read, choosing one alignment randomly in case of a tie. Peaks were called using MACS2 callpeak (v2.2.6) on default settings using MACS2’s shifting model (Zhang et al. 2008) (https://github.com/macs3-project/MACS). Broad peaks were called at a FDR of 5%, while narrow peaks were called at a FDR of 1%. BigWig files for peak’s visualization were obtained with MACS2 bdgcmp tool and UCSC bedGraphToBigWig. For H3K27me3, which occurs in large domains, enriched regions were identified with hiddenDomains (Starmer and Magnuson 2016). Paired-end long-ChIP reads were generated as described above. Illumina adapters were removed using Trimmomatic (parameters SLIDINGWINDOW:4:30 MINLEN:50). Reads were mapped using both paired-end BWA mem and single-end Bowtie allowing for multiple mappings (parameters -a -n -S -e 200 -m 99). For single-end alignments, forward and reverse reads were concatenated into a single file and properly renamed to secure unique reads IDs. Reads aligned with BWA were filtered by MAPQ ≥ 20 while reads with multiple mappings aligned with Bowtie were allocated with CSEM and most likely alignments were selected with the custom script. Duplicates and secondary alignments were removed as explained above. Peaks were called using MACS2 with identical parameters used for short-reads, adding the BAMPE option in the case of paired-end reads aligned with BWA mem. Sets of peaks were compared between analysis methods using HOMER mergePeaks (parameters: “-d given”) (Heinz et al. 2010) and a unidirectional correlation metric derived from IntervalStats using peaks with an overlap *p*-value below 0.05 (Chikina and Troyanskaya 2012).

For depletion analyses, SD coordinates were directly downloaded from UCSC Table Browser and HSD coordinates were obtained by filtering alignments with sequence identity over 98% in the fracMatch column, converting them to BED format and merging overlapping entries using bedtools merge. The number of peaks and bases under peaks on each region of interest were obtained with bedtools intersect. To obtain depletion statistics, 1000 regions of the same size as SD and HSD were randomly sampled from the human genome GRCh38. Empirical *p*-values of depletion tests were calculated as *p*-value = (M+1)/(N+1), where M is the number of iterations less than the observed value and N is the number of iterations.

Additionally, mapping quality scores (MAPQ) distributions for H3K27ac following a similar approach as explained before, but using BWA aln and BWA mem for short and long ChIP-seq reads respectively, after PCR duplicates and secondary alignments removal. Posterior probabilities distributions for H3K27ac were examined using the output of CSEM after selecting the most likely alignment with the custom script. Entries in unique space were subsampled to 10 million and plots were obtained with the geom_density function in ggplot R package.

All ChIP-seq analyses are available as a TrackHub for the UCSC Genome Browser (https://bioshare.bioinformatics.ucdavis.edu/bioshare/download/cpqqdfge5lfvovq/hsd_noncoding/hub.txt). Further, the bioinformatic pipeline is freely available for use in Snakemake format (https://github.com/mydennislab/snake-chipseq), allowing the analysis to be replicated in any cell or tissue type of interest.

### Paralog-specific validation of RNA expression and ChIP data

Following published protocols (Integrated DNA Technologies), we used the rhAMP assay in 10 µl total reaction volumes to quantify abundance of PSVs (for all assays except *ARHGAP11* expression, the fluorophores FAM=A paralog and VIC=B paralog) as a proxy for paralog-specific expression (RNA) and enrichment (ChIP) (Table S7). We used 10 ng total of RNA converted to cDNA to validate gene expression for duplicated gene families *ARHGAP11, ROCK1*, and *DUSP22*. We calculated dCt of cDNA and gDNA as Ct_FAM_-Ct_VIC_ and ddCt as dCt_cDNA_-dCt_gDNA_ from the same cell line. We calculated dCt of the input and ChIP-enriched library as Ct_FAM_-Ct_VIC_ and ddCt as dCt_ChIP_-dCt_input_ from the same cell line. For both expression and ChIP analyses, the ratio of abundance of the B to the A paralog is 2^ddCt^.

### ChromHMM annotations

We generated ChromHMM (version 1.19) (Ernst and Kellis 2012) models separately for ENCODE short-read data and long ChIP after multi-mapping and CSEM allocation, using active chromatin histone modifications (H3K4me3, H3K4me1, and H3K27ac). States corresponding to active transcription start sites and active enhancers were identified manually (Ernst and Kellis 2017). In the ENCODE analysis, promoters were assigned to state 1, which corresponded to active transcription start sites, and enhancers were assigned to state 8, which corresponded to active enhancers (Figure S9A). Similarly, in the long ChIP analysis, promoters were assigned to state 3 and active enhancers were assigned to states 6; state 4 was considered to be an additional enhancer state lacking enrichment in H3K4me1 (Ernst and Kellis 2017) (Figure S9B). Together, these sets of elements were defined as cCREs.

### Luciferase reporter assays

Promoters of highly and differentially expressed HSD gene families (*ARHGAP11, NCF1*, and *DUSP22*) were chosen for screening in a reporter assay. Fragments containing the TSS and spanning ∼1 kb were amplified with KpnI and SacI restriction sites included in primers (Table S7) and cloned into the luciferase reporter vector pGL3-basic (Promega). Candidate enhancers within 50 kb of genes bodies were selected based on the presence of ChromHMM CREs in the re-analyzed data from human LCLs. Target regions were a maximum size of 5 kb, and peaks larger than this were tiled with multiple targets. Gateway homology arms were added to primers in accordance with the manual (ThermoFisher), and PCR products were cloned into the entry vector pDONR221 (ThermoFisher 12536017). Expression clones for luciferase assays were generated by cloning pDONR221 inserts into the luciferase reporter pE1B (Antonellis et al. 2008) with the Gateway system.

Constructs were co-transfected (ThermoFisher Lipofectamine 3000) in equimolar amounts with 50 ng of the control plasmid pRL-TK (Renilla luciferase) into HeLa cells in 96-well plates. Cells were at 70-90% confluence at the time of transfection. Luciferase assays were performed with the Dual-Luciferase Reporter Assay System (Promega E1910). 48 hours post-transfection, cells were washed with PBS, and lysed with Passive Lysis Buffer for at least 15 min sharking at 500 rpm. Lysates were stored at -80C. For LCLs, cells were split 48 and 24 hours pre-transfection to ensure active division. Cells were counted, washed in PBS, and resuspended such that each transfection contained 12.5×10^6^ cells, 6.25 ug of test construct, and equimolar pRL-TK in RPMI. Cells were electroporated using the Neon Transfection System in accordance with previously published work (Tewhey et al. 2018) and recovered at a density of 3×10^6^ cells/mL in pre-warmed RPMI including 15% FBS without antibiotics. Transfection efficiencies of ∼15% were achieved. To perform luciferase assays, ∼5×10^5^ cells were pipetted into each well of a 96-well plate, washed with PBS, and lysed with Passive Lysis Buffer as described for HeLa. Luminescence measurements were performed according to the manufacturer’s instructions using a Tecan Infinite or Tecan Spark plate reader with injectors.

### Transcription factors binding motifs

Alignments of cloned sequences were scanned for HOmo sapiens COmprehensive MOdel COllection (HOCOMOCO) v11 (Kulakovskiy et al. 2018) TFBS motifs using FIMO (Grant et al. 2011). HOCOMOCO motifs were limited to transcription factors expressed above 1 TPM in >75% of ENCODE mRNA-seq libraries generated for GM12878 (ENCSR077AZT, ENCLB555AQG, ENCLB555AQH, ENCLB555ANP, ENCLB555ALI, ENCLB555ANM, ENCLB555ANN, ENCLB037ZZZ, ENCLB038ZZZ, ENCLB043ZZZ, ENCLB044ZZZ, ENCLB041ZZZ, ENCLB042ZZZ, ENCLB045ZZZ, ENCLB046ZZZ, ENCLB700LMU, ENCLB150CGC). Significant matches above a 5% FDR were retained for the analysis. TFBSs were compared across homologous sequences to identify putative paralog-specific gains and losses of binding sites.

## ACKNOWLEDGMENTS

We thank the many groups/consortia that have made their data publicly available, including Dr. Yoav Gilad, the 1000 Genomes Project, GTEx, and ENCODE for which this research would not be possible without its use. In particular, we acknowledge the labs of Dr. Bradley Bernstein, Dr. Ali Mortazavi, Dr. Barbara Wold, Dr. Thomas Gingeras, and Dr. Brenton Graveley, which generated the ENCODE data used in this publication. We also thank Dr. Colin Kern for valuable advice concerning ChIP-seq analysis, Dr. Anthony Antonellis for sharing the pE1B enhancer reporter Gateway plasmid, and Drs. Gerald Quon and Siobhan Brady for constructive feedback on the manuscript. This work was supported by the National Human Genome Research Institute (F31HG011205 to C.S.), National Institute of Neurological Disorders and Stroke (R00NS083627 to M.Y.D.), and the Office of the Director and National Institute of Mental Health (DP2 OD025824 to M.Y.D.) at the National Institutes of Health (NIH). Statistical analysis advice was provided by Dr. Blythe Durbin-Johnson through the MIND Institute Intellectual and Developmental Disability Research Center, funded by the NIH National Institute of Child Health and Human Development (U54 HD079125). Additionally, M.Y.D. is supported as a Sloan fellow (FG-2016-6814), P.C.M as an NIH National Institute of Mental Health T32 UC Davis Autism Research Training Program fellow (T32MH073124-17), D.C.S. as a Fulbright fellow, and J.R. as an NIH National Institute of General Medical Sciences UC Davis Postbaccalaureate Research Education Program fellow (R25GM116690).

## DATA ACCESS

Large-insert ChIP-sequencing data generated for this study are available from the European Nucleotide Archive under the accession PRJEB40356.

## References

Agarwal V, Bell GW, Nam J-W, Bartel DP. 2015. Predicting effective microRNA target sites in mammalian mRNAs. eLife [Internet] 4. Available from: http://dx.doi.org/10.7554/elife.05005

Altschul SF, Gish W, Miller W, Myers EW, Lipman DJ. 1990. Basic local alignment search tool. J. Mol. Biol. 215:403–410.

Anders S, Pyl PT, Huber W. 2015. HTSeq--a Python framework to work with high-throughput sequencing data. Bioinformatics 31:166–169.

Anon. 2013. Potential role of human-specific genes, human-specific microRNAs and human-specific non-coding regulatory RNAs in the pathogenesis of Systemic Sclerosis and Sjögren’s Syndrome. Autoimmun. Rev. 12:1046–1051.

Anon. 2017. Are humans prone to autoimmunity? Implications from evolutionary changes in hominin sialic acid biology. J. Autoimmun. 83:134–142.

Antonacci F, Dennis MY, Huddleston J, Sudmant PH, Steinberg KM, Rosenfeld JA, Miroballo M, Graves TA, Vives L, Malig M, et al. 2014. Palindromic GOLGA8 core duplicons promote chromosome 15q13.3 microdeletion and evolutionary instability. Nature Genetics [Internet] 46:1293–1302. Available from: http://dx.doi.org/10.1038/ng.3120

Antonellis A, Huynh JL, Lee-Lin S-Q, Vinton RM, Renaud G, Loftus SK, Elliot G, Wolfsberg TG, Green ED, McCallion AS, et al. 2008. Identification of neural crest and glial enhancers at the mouse Sox10 locus through transgenesis in zebrafish. PLoS Genet. 4:e1000174.

Bailey JA. 2002. Recent Segmental Duplications in the Human Genome. Science [Internet] 297:1003– 1007. Available from: http://dx.doi.org/10.1126/science.1072047

Barreiro LB, Marioni JC, Blekhman R, Stephens M, Gilad Y. 2010. Functional Comparison of Innate Immune Signaling Pathways in Primates. PLoS Genet. 6:e1001249.

Blake LE, Roux J, Hernando-Herraez I, Banovich NE, Perez RG, Hsiao CJ, Eres I, Cuevas C, Marques-Bonet T, Gilad Y. 2020. A comparison of gene expression and DNA methylation patterns across tissues and species. Genome Res. 30:250–262.

Bolger AM, Lohse M, Usadel B. 2014. Trimmomatic: a flexible trimmer for Illumina sequence data. Bioinformatics 30:2114–2120.

Braasch I, Bobe J, Guiguen Y, Postlethwait JH. 2018. Reply to: “Subfunctionalization versus neofunctionalization after whole-genome duplication.” Nat. Genet. 50:910–911.

Braasch I, Gehrke AR, Smith JJ, Kawasaki K, Manousaki T, Pasquier J, Amores A, Desvignes T, Batzel P, Catchen J, et al. 2016. The spotted gar genome illuminates vertebrate evolution and facilitates human-teleost comparisons. Nature Genetics [Internet] 48:427–437. Available from: http://dx.doi.org/10.1038/ng.3526

Charrier C, Joshi K, Coutinho-Budd J, Kim J-E, Lambert N, de Marchena J, Jin W-L, Vanderhaeghen P, Ghosh A, Sassa T, et al. 2012. Inhibition of SRGAP2 Function by Its Human-Specific Paralogs Induces Neoteny during Spine Maturation. Cell [Internet] 149:923–935. Available from: http://dx.doi.org/10.1016/j.cell.2012.03.034

Chikina MD, Troyanskaya OG. 2012. An effective statistical evaluation of ChIPseq dataset similarity. Bioinformatics 28:607–613.

Chung D, Kuan PF, Li B, Sanalkumar R, Liang K, Bresnick EH, Dewey C, Keleş S. 2011. Discovering transcription factor binding sites in highly repetitive regions of genomes with multi-read analysis of ChIP-Seq data. PLoS Comput. Biol. 7:e1002111.

Consortium TEP, The ENCODE Project Consortium. 2012. An integrated encyclopedia of DNA elements in the human genome. Nature [Internet] 489:57–74. Available from: http://dx.doi.org/10.1038/nature11247

Davis CA, Hitz BC, Sloan CA, Chan ET, Davidson JM, Gabdank I, Hilton JA, Jain K, Baymuradov UK, Narayanan AK, et al. 2018. The Encyclopedia of DNA elements (ENCODE): data portal update.

Nucleic Acids Research [Internet] 46:D794–D801. Available from: http://dx.doi.org/10.1093/nar/gkx1081

Dennis MY, Eichler EE. 2016. Human adaptation and evolution by segmental duplication. Current Opinion in Genetics & Development [Internet] 41:44–52. Available from: http://dx.doi.org/10.1016/j.gde.2016.08.001

Dennis MY, Harshman L, Nelson BJ, Penn O, Cantsilieris S, Huddleston J, Antonacci F, Penewit K, Denman L, Raja A, et al. 2017. The evolution and population diversity of human-specific segmental duplications. Nat Ecol Evol 1:69.

Dennis MY, Nuttle X, Sudmant PH, Antonacci F, Graves TA, Nefedov M, Rosenfeld JA, Sajjadian S, Malig M, Kotkiewicz H, et al. 2012. Evolution of Human-Specific Neural SRGAP2 Genes by Incomplete Segmental Duplication. Cell [Internet] 149:912–922. Available from: http://dx.doi.org/10.1016/j.cell.2012.03.033

Dougherty ML, Nuttle X, Penn O, Nelson BJ, Huddleston J, Baker C, Harshman L, Duyzend MH, Ventura M, Antonacci F, et al. 2017. The birth of a human-specific neural gene by incomplete duplication and gene fusion. Genome Biol. 18:49.

Dougherty ML, Underwood JG, Nelson BJ, Tseng E, Munson KM, Penn O, Nowakowski TJ, Pollen AA, Eichler EE. 2018. Transcriptional fates of human-specific segmental duplications in brain. Genome Research [Internet] 28:1566–1576. Available from: http://dx.doi.org/10.1101/gr.237610.118

Dumont BL. 2015. Interlocus gene conversion explains at least 2.7 % of single nucleotide variants in human segmental duplications. BMC Genomics [Internet] 16. Available from: http://dx.doi.org/10.1186/s12864-015-1681-3

Ebbert MTW, Jensen TD, Jansen-West K, Sens JP, Reddy JS, Ridge PG, Kauwe JSK, Belzil V, Pregent L, Carrasquillo MM, et al. 2019. Systematic analysis of dark and camouflaged genes reveals disease-relevant genes hiding in plain sight. Genome Biol. 20:97.

Ernst J, Kellis M. 2012. ChromHMM: automating chromatin-state discovery and characterization. Nature Methods [Internet] 9:215–216. Available from: http://dx.doi.org/10.1038/nmeth.1906

Ernst J, Kellis M. 2017. Chromatin-state discovery and genome annotation with ChromHMM. Nat. Protoc. 12:2478–2492.

Fairley S, Lowy-Gallego E, Perry E, Flicek P. 2020. The International Genome Sample Resource (IGSR) collection of open human genomic variation resources. Nucleic Acids Res. 48:D941–D947.

Fiddes IT, Lodewijk GA, Mooring M, Bosworth CM, Ewing AD, Mantalas GL, Novak AM, van den Bout A, Bishara A, Rosenkrantz JL, et al. 2018. Human-Specific NOTCH2NL Genes Affect Notch Signaling and Cortical Neurogenesis. Cell [Internet] 173:1356–1369.e22. Available from: http://dx.doi.org/10.1016/j.cell.2018.03.051

Florio M, Albert M, Taverna E, Namba T, Brandl H, Lewitus E, Haffner C, Sykes A, Wong FK, Peters J, et al. 2015. Human-specific gene ARHGAP11B promotes basal progenitor amplification and neocortex expansion. Science [Internet] 347:1465–1470. Available from: http://dx.doi.org/10.1126/science.aaa1975

Genomes Project, Consortium, Auton A, Brooks LD, Durbin RM, Garrison EP, Kang HM, Korbel JO, Marchini JL, McCarthy S, McVean GA, et al. 2015. A global reference for human genetic variation. Nature 526:68–74.

Giannuzzi G, Migliavacca E, Reymond A. 2014. Novel H3K4me3 marks are enriched at human- and chimpanzee-specific cytogenetic structures. Genome Research [Internet] 24:1455–1468. Available from: http://dx.doi.org/10.1101/gr.167742.113

Grant CE, Bailey TL, Noble WS. 2011. FIMO: scanning for occurrences of a given motif. Bioinformatics [Internet] 27:1017–1018. Available from: http://dx.doi.org/10.1093/bioinformatics/btr064

Hartasánchez DA, Brasó-Vives M, Heredia-Genestar JM, Pybus M, Navarro A. 2018. Effect of Collapsed Duplications on Diversity Estimates: What to Expect. Genome Biol. Evol. 10:2899–2905.

Heide M, Haffner C, Murayama A, Kurotaki Y, Shinohara H, Okano H, Sasaki E, Huttner WB. 2020. Human-specific ARHGAP11B increases size and folding of primate neocortex in the fetal marmoset. Science [Internet]:eabb2401. Available from: http://dx.doi.org/10.1126/science.abb2401

Heinz S, Benner C, Spann N, Bertolino E, Lin YC, Laslo P, Cheng JX, Murre C, Singh H, Glass CK. 2010. Simple Combinations of Lineage-Determining Transcription Factors Prime cis-Regulatory Elements Required for Macrophage and B Cell Identities. Molecular Cell [Internet] 38:576–589. Available from: http://dx.doi.org/10.1016/j.molcel.2010.05.004

Hultqvist M, Olofsson P, Holmberg J, Backstrom BT, Tordsson J, Holmdahl R. 2004. Enhanced autoimmunity, arthritis, and encephalomyelitis in mice with a reduced oxidative burst due to a mutation in the Ncf1 gene. Proceedings of the National Academy of Sciences [Internet] 101:12646–12651. Available from: http://dx.doi.org/10.1073/pnas.0403831101

Ishiura H, Shibata S, Yoshimura J, Suzuki Y, Qu W, Doi K, Asem Almansour M, Kikuchi JK, Taira M, Mitsui J, et al. 2019. Noncoding CGG repeat expansions in neuronal intranuclear inclusion disease, oculopharyngodistal myopathy and an overlapping disease. Nature Genetics [Internet] 51:1222– 1232. Available from: http://dx.doi.org/10.1038/s41588-019-0458-z

Juric I, Yu M, Abnousi A, Raviram R, Fang R, Zhao Y, Zhang Y, Qiu Y, Yang Y, Li Y, et al. 2019. MAPS: Model-based analysis of long-range chromatin interactions from PLAC-seq and HiChIP experiments. PLoS Comput. Biol. 15:e1006982.

Kalebic N, Gilardi C, Albert M, Namba T, Long KR, Kostic M, Langen B, Huttner WB. 2018. Human-specific ARHGAP11B induces hallmarks of neocortical expansion in developing ferret neocortex. eLife [Internet] 7. vailable from: http://dx.doi.org/10.7554/elife.41241

Kassahn KS, Dang VT, Wilkins SJ, Perkins AC, Ragan MA. 2009. Evolution of gene function and regulatory control after whole-genome duplication: comparative analyses in vertebrates. Genome Res. 19:1404–1418.

Kent WJ. 2002. BLAT--the BLAST-like alignment tool. Genome Res. 12:656–664.

Khan Z, Ford MJ, Cusanovich DA, Mitrano A, Pritchard JK, Gilad Y. 2013. Primate transcript and protein expression levels evolve under compensatory selection pressures. Science 342:1100–1104.

Kronenberg ZN, Fiddes IT, Gordon D, Murali S, Cantsilieris S, Meyerson OS, Underwood JG, Nelson BJ, Chaisson MJP, Dougherty ML, et al. 2018. High-resolution comparative analysis of great ape genomes. Science [Internet] 360. Available from: http://dx.doi.org/10.1126/science.aar6343

Kulakovskiy IV, Vorontsov IE, Yevshin IS, Sharipov RN, Fedorova AD, Rumynskiy EI, Medvedeva YA, Magana-Mora A, Bajic VB, Papatsenko DA, et al. 2018. HOCOMOCO: towards a complete collection of transcription factor binding models for human and mouse via large-scale ChIP-Seq analysis. Nucleic Acids Res. 46:D252–D259.

Landt SG, Marinov GK, Kundaje A, Kheradpour P, Pauli F, Batzoglou S, Bernstein BE, Bickel P, Brown JB, Cayting P, et al. 2012. ChIP-seq guidelines and practices of the ENCODE and modENCODE consortia. Genome Res. 22:1813–1831.

Langmead B, Trapnell C, Pop M, Salzberg SL. 2009. Ultrafast and memory-efficient alignment of short DNA sequences to the human genome. Genome Biol. 10:R25.

Lan X, Pritchard JK. 2016. Coregulation of tandem duplicate genes slows evolution of subfunctionalization in mammals. Science [Internet] 352:1009–1013. Available from: http://dx.doi.org/10.1126/science.aad8411

Lappalainen T, The Geuvadis Consortium, Sammeth M, Friedländer MR, ‘t PA Monlong J, Rivas MA, Gonzàlez-Porta M, Kurbatova N, Griebel T, et al. 2013. Transcriptome and genome sequencing uncovers functional variation in humans. Nature [Internet] 501:506–511. Available from: http://dx.doi.org/10.1038/nature12531

Law CW, Chen Y, Shi W, Smyth GK. 2014. voom: precision weights unlock linear model analysis tools for RNA-seq read counts. Genome Biology [Internet] 15:R29. Available from: http://dx.doi.org/10.1186/gb-2014-15-2-r29

Lien S, Koop BF, Sandve SR, Miller JR, Kent MP, Nome T, Hvidsten TR, Leong JS, Minkley DR, Zimin A, et al. 2016. The Atlantic salmon genome provides insights into rediploidization. Nature [Internet] 533:200–205. Available from: http://dx.doi.org/10.1038/nature17164

Li J-P, Yang C-Y, Chuang H-C, Lan J-L, Chen D-Y, Chen Y-M, Wang X, Chen AJ, Belmont JW, Tan T-H. 2014. The phosphatase JKAP/DUSP22 inhibits T-cell receptor signalling and autoimmunity by inactivating Lck. Nature Communications [Internet] 5. Available from: http://dx.doi.org/10.1038/ncomms4618

Lynch M. 2000. The Evolutionary Fate and Consequences of Duplicate Genes. Science [Internet] 290:1151–1155. Available from: http://dx.doi.org/10.1126/science.290.5494.1151

Marchetto MC, Hrvoj-Mihic B, Kerman BE, Yu DX, Vadodaria KC, Linker SB, Narvaiza I, Santos R, Denli AM, Mendes APD, et al. 2019. Species-specific maturation profiles of human, chimpanzee and bonobo neural cells. eLife [Internet] 8. Available from: http://dx.doi.org/10.7554/elife.37527

McVicker G, van de Geijn B, Degner JF, Cain CE, Banovich NE, Raj A, Lewellen N, Myrthil M, Gilad Y, Pritchard JK. 2013. Identification of Genetic Variants That Affect Histone Modifications in Human Cells. Science [Internet] 342:747–749. Available from: http://dx.doi.org/10.1126/science.1242429

Mifsud B, Tavares-Cadete F, Young AN, Sugar R, Schoenfelder S, Ferreira L, Wingett SW, Andrews S, Grey W, Ewels PA, et al. 2015. Mapping long-range promoter contacts in human cells with high-resolution capture Hi-C. Nat. Genet. 47:598–606.

Mumbach MR, Satpathy AT, Boyle EA, Dai C, Gowen BG, Cho SW, Nguyen ML, Rubin AJ, Granja JM, Kazane KR, et al. 2017. Enhancer connectome in primary human cells identifies target genes of disease-associated DNA elements. Nat. Genet. 49:1602–1612.

Nguyen LS, Jolly L, Shoubridge C, Chan WK, Huang L, Laumonnier F, Raynaud M, Hackett A, Field M, Rodriguez J, et al. 2012. Transcriptome profiling of UPF3B/NMD-deficient lymphoblastoid cells from patients with various forms of intellectual disability. Molecular Psychiatry [Internet] 17:1103– 1115. Available from: http://dx.doi.org/10.1038/mp.2011.163

Noensie EN, Dietz HC. 2001. A strategy for disease gene identification through nonsense-mediated mRNA decay inhibition. Nat. Biotechnol. 19:434–439.

O’Bleness M, Searles VB, Dickens CM, Astling D, Albracht D, Mak AC, Lai YY, Lin C, Chu C, Graves T, et al. 2014. Finished sequence and assembly of the DUF1220-rich 1q21 region using a haploid human genome. BMC Genomics 15:387.

O’Geen H, Bates SL, Carter SS, Nisson KA, Halmai J, Fink KD, Rhie SK, Farnham PJ, Segal DJ. 2019. Ezh2-dCas9 and KRAB-dCas9 enable engineering of epigenetic memory in a context-dependent manner. Epigenetics Chromatin 12:26.

Ohno S. 1970. Evolution by Gene Duplication. Available from: http://dx.doi.org/10.1007/978-3-642-86659-386659-386659-386659-3

Patro R, Duggal G, Love MI, Irizarry RA, Kingsford C. 2017. Salmon provides fast and bias-aware quantification of transcript expression. Nature Methods [Internet] 14:417–419. Available from: http://dx.doi.org/10.1038/nmeth.4197

Pavlovic BJ, Blake LE, Roux J, Chavarria C, Gilad Y. 2018. A Comparative Assessment of Human and Chimpanzee iPSC-derived Cardiomyocytes with Primary Heart Tissues. Sci. Rep. 8:15312.

Pickrell JK, Marioni JC, Pai AA, Degner JF, Engelhardt BE, Nkadori E, Veyrieras J-B, Stephens M, Gilad Y, Pritchard JK. 2010. Understanding mechanisms underlying human gene expression variation with RNA sequencing. Nature 464:768–772.

Prado-Martinez J, Sudmant PH, Kidd JM, Li H, Kelley JL, Lorente-Galdos B, Veeramah KR, Woerner AE, O’Connor TD, Santpere G, et al. 2013. Great ape genetic diversity and population history. Nature 499:471–475.

Qian W, Liao B-Y, Chang AY-F, Zhang J. 2010. Maintenance of duplicate genes and their functional redundancy by reduced expression. Trends in Genetics [Internet] 26:425–430. Available from: http://dx.doi.org/10.1016/j.tig.2010.07.002

Ritchie ME, Phipson B, Wu D, Hu Y, Law CW, Shi W, Smyth GK. 2015. limma powers differential expression analyses for RNA-sequencing and microarray studies. Nucleic Acids Res. 43:e47.

Rodin SN, Parkhomchuk DV, Riggs AD. 2005. Epigenetic changes and repositioning determine the evolutionary fate of duplicated genes. Biochemistry 70:559–567.

Rodin SN, Riggs AD. 2003. Epigenetic silencing may aid evolution by gene duplication. J. Mol. Evol. 56:718–729.

Sandve SR, Rohlfs RV, Hvidsten TR. 2018. Subfunctionalization versus neofunctionalization after whole-genome duplication. Nature Genetics [Internet] 50:908–909. Available from: http://dx.doi.org/10.1038/s41588-018-0162-4

Shen F, Kidd JM. 2020. Rapid, Paralog-Sensitive CNV Analysis of 2457 Human Genomes Using QuicK- mer2. Genes [Internet] 11. Available from: http://dx.doi.org/10.3390/genes11020141

Soneson C, Love MI, Robinson MD. 2015. Differential analyses for RNA-seq: transcript-level estimates improve gene-level inferences. F1000Res. 4:1521.

Starmer J, Magnuson T. 2016. Detecting broad domains and narrow peaks in ChIP-seq data with hiddenDomains. BMC Bioinformatics 17:144.

Steinberg KM, Antonacci F, Sudmant PH, Kidd JM, Campbell CD, Vives L, Malig M, Scheinfeldt L, Beggs W, Ibrahim M, et al. 2012. Structural diversity and African origin of the 17q21.31 inversion polymorphism. Nature Genetics [Internet] 44:872–880. Available from: http://dx.doi.org/10.1038/ng.2335

Stranger BE, Forrest MS, Dunning M, Ingle CE, Beazley C, Thorne N, Redon R, Bird CP, de Grassi A, Lee C, et al. 2007. Relative impact of nucleotide and copy number variation on gene expression phenotypes. Science 315:848–853.

Suzuki IK, Gacquer D, Van Heurck R, Kumar D, Wojno M, Bilheu A, Herpoel A, Lambert N, Cheron J, Polleux F, et al. 2018. Human-Specific NOTCH2NL Genes Expand Cortical Neurogenesis through Delta/Notch Regulation. Cell [Internet] 173:1370–1384.e16. Available from: http://dx.doi.org/10.1016/j.cell.2018.03.067

Tewhey R, Kotliar D, Park DS, Liu B, Winnicki S, Reilly SK, Andersen KG, Mikkelsen TS, Lander ES, Schaffner SF, et al. 2018. Direct Identification of Hundreds of Expression-Modulating Variants using a Multiplexed Reporter Assay. Cell 172:1132–1134.

Tvrdik P, Capecchi MR. 2006. Reversal of Hox1 gene subfunctionalization in the mouse. Dev. Cell 11:239–250.

Varadharajan S, Sandve SR, Gillard GB, Tørresen OK, Mulugeta TD, Hvidsten TR, Lien S, Vøllestad LA, Jentoft S, Nederbragt AJ, et al. 2018. The Grayling Genome Reveals Selection on Gene Expression Regulation after Whole-Genome Duplication. Genome Biology and Evolution [Internet] 10:2785–2800. Available from: http://dx.doi.org/10.1093/gbe/evy201

Wen X, Luca F, Pique-Regi R. 2015. Cross-Population Joint Analysis of eQTLs: Fine Mapping and Functional Annotation. PLOS Genetics [Internet] 11:e1005176. Available from: http://dx.doi.org/10.1371/journal.pgen.1005176

Zhang J. 2003. Evolution by gene duplication: an update. Trends in Ecology & Evolution [Internet] 18:292–298. Available from: http://dx.doi.org/10.1016/s0169-5347(03)00033-8

Zhang Y, Liu T, Meyer CA, Eeckhoute J, Johnson DS, Bernstein BE, Nusbaum C, Myers RM, Brown M, Li W, et al. 2008. Model-based analysis of ChIP-Seq (MACS). Genome Biol. 9:R137.

Zhao J, Ma J, Deng Y, Kelly JA, Kim K, Bang S-Y, Lee H-S, Li Q-Z, Wakeland EK, Qiu R, et al. 2017. A missense variant in NCF1 is associated with susceptibility to multiple autoimmune diseases. Nature Genetics [Internet] 49:433–437. Available from: http://dx.doi.org/10.1038/ng.3782

Zheng Y, Ay F, Keles S. 2019. Generative modeling of multi-mapping reads with mHi-C advances analysis of Hi-C studies. Elife [Internet] 8. Available from: http://dx.doi.org/10.7554/eLife.38070

