## Supplementary Figures for "Diverse molecular mechanisms contribute to differential expression of human duplicated genes"

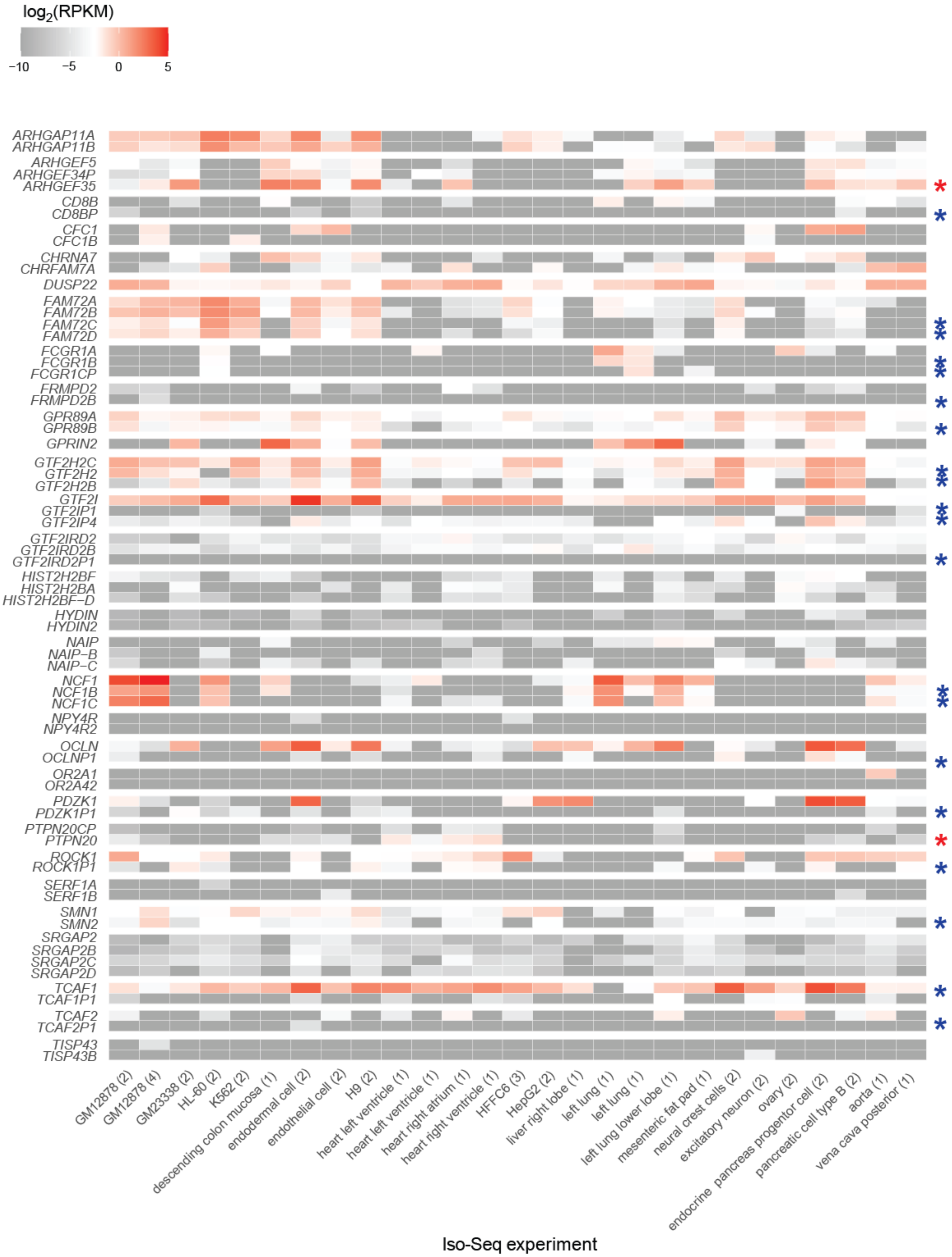


**Figure S1. HSD gene expression in Iso-Seq datasets.** For each available ENCODE Iso-Seq experiment (columns; number of technical replicates indicated in parentheses), HSD gene expression was calculated in reads per kilobase per million mapped reads (RPKM) using only paralogous regions (i.e., excluding truncated portions and novel portions of genes). For derived genes, $\log_2$(RPKM) were compared to the ancestral gene with a Wilcoxon signed-rank test. Significant differences (Benjamini-Hochberg adjusted $p < 0.05$) are indicated with an asterisk (blue for lower expression; red for higher). *DUSP22* and *GPRIN2* were not tested for differential expression because their derived genes are missing from GRCh38.

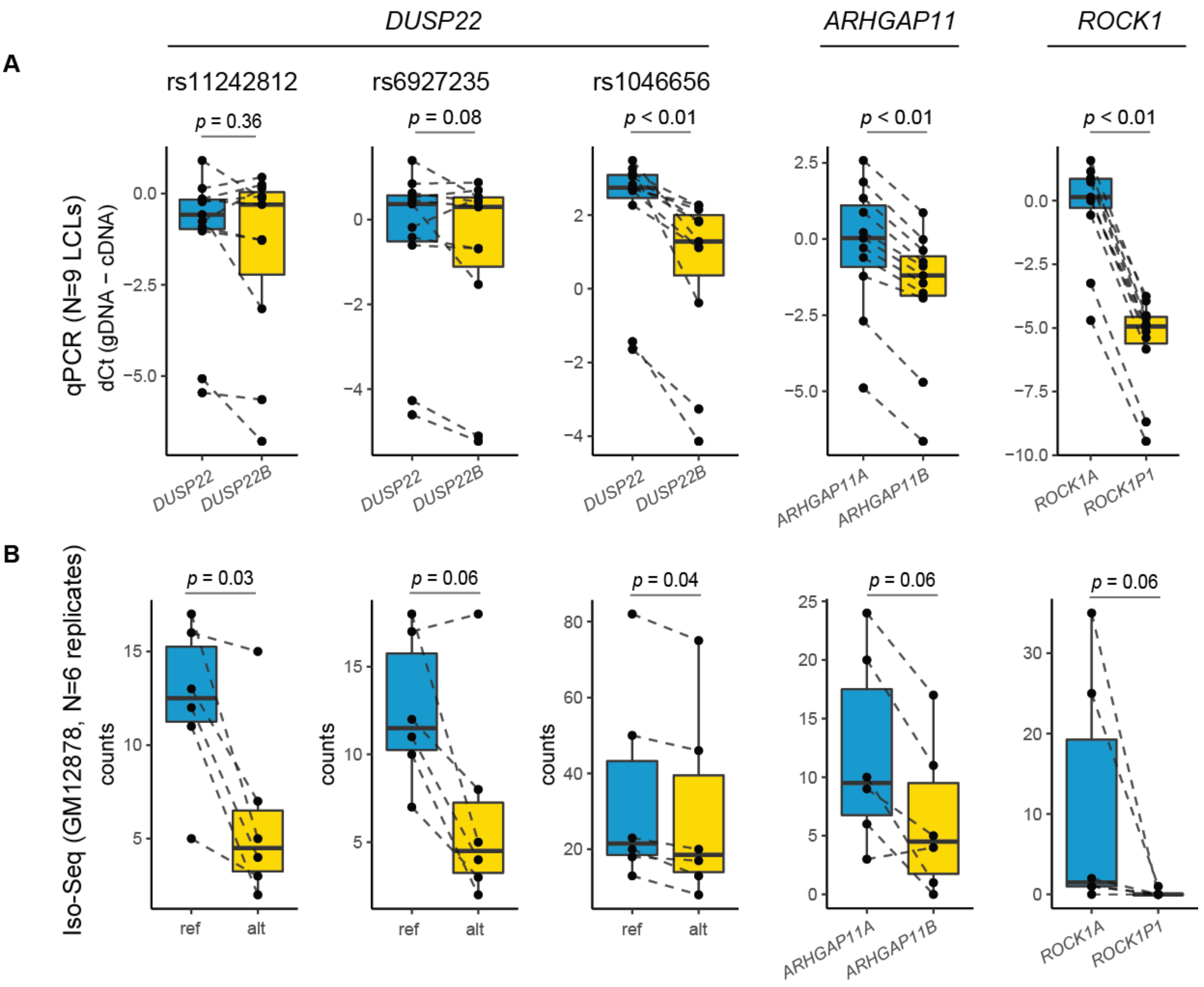


**Figure S2. Validation of short-read RNA-seq quantification. (A)** qPCR was conducted on genomic DNA (gDNA) and complementary DNA (cDNA) extracted from nine human LCLs to quantify expression differences at paralog-specific variants (PSVs). Because *DUSP22B* is missing from the reference, *DUSP22* PSVs are annotated as SNPs, where the alternate allele corresponds to a known *DUSP22B* substitution. The difference of cycle threshold (dCt) between gDNA and cDNA samples was calculated for each LCL and compared between paralogs with a paired Wilcoxon signed-rank test. Each point represents the mean of three technical replicates. **(B)** Read counts from six replicates of GM12878 Iso-Seq experiments (ENCODE) at *DUSP22* PSVs (raw alignments prior to sequence correction) or *ARHGAP11*/*ROCK1* paralogous regions (filtered alignments). Differences were quantified with a paired Wilcoxon signed-rank test.

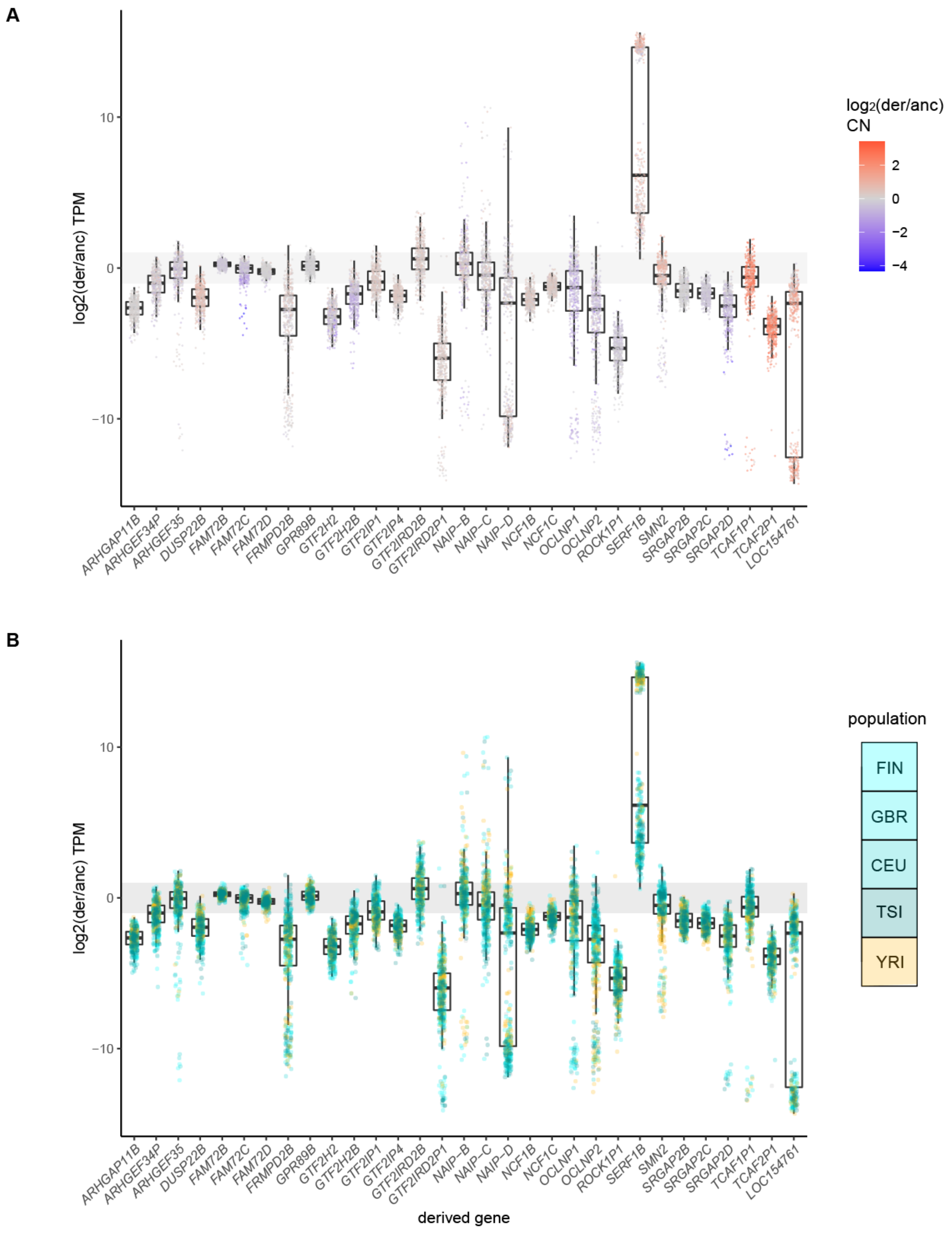


**Figure S3. Expression divergence of derived HSD genes.** Expression divergence of derived genes from families with at least one LCL-expressed paralog is plotted as the $log_2$ ratio of median derived and ancestral TPM expression. Each point represents a different LCL from the Geuvadis consortium (total N=445). The gray bar indicates a two-fold expression difference. Colors represent **(A)** relative CN and **(B)** LCL source population.

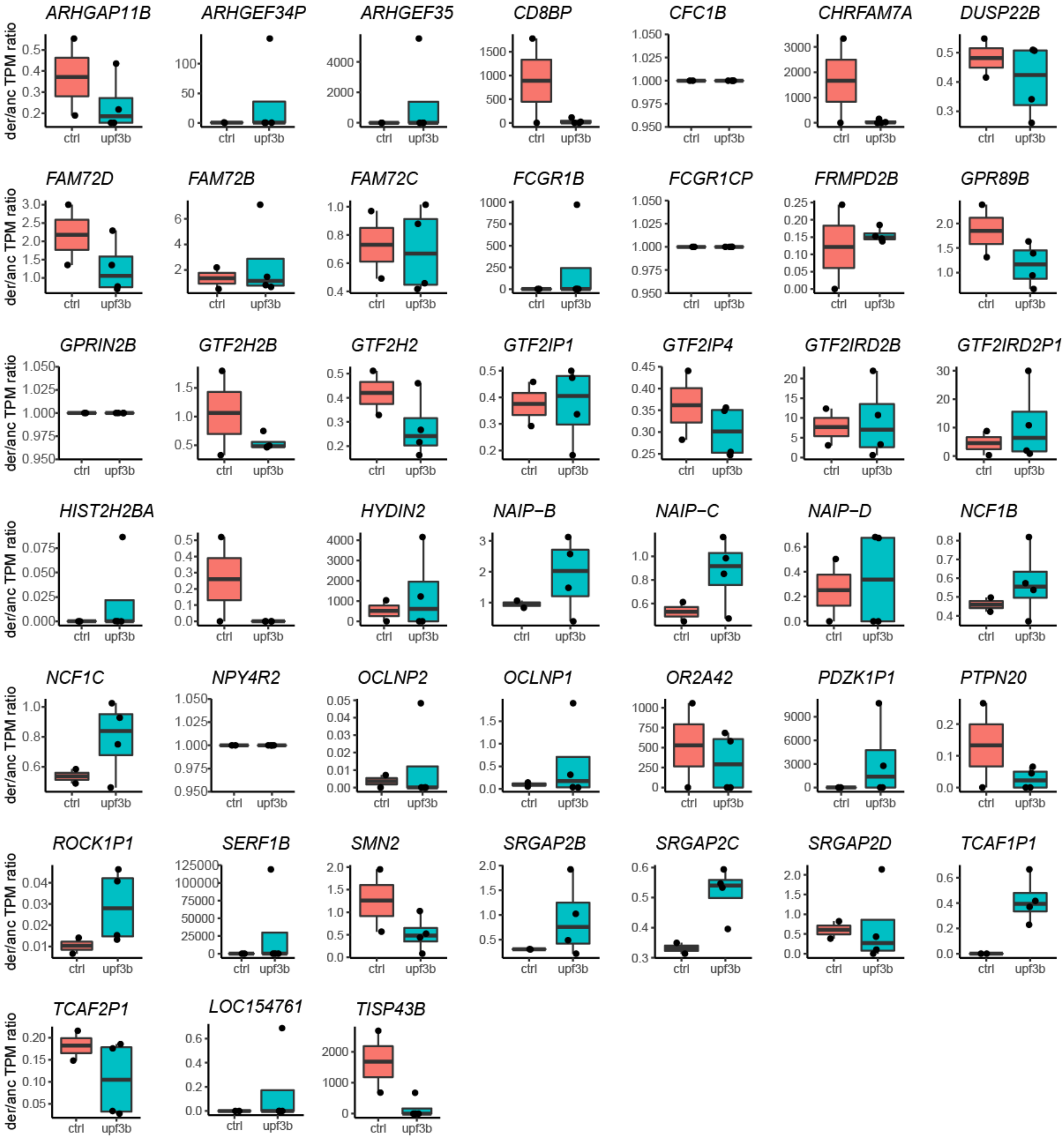


**Figure S4. Nonsense-mediated decay of HSD transcripts.** Derived/ancestral TPM ratios of HSD genes, calculated from RNA-seq of control (ctrl) and NMD-deficient cells (upf3b) (Nguyen et al. 2012). No differences were determined to be significant by differential expression analysis (limma-voom).

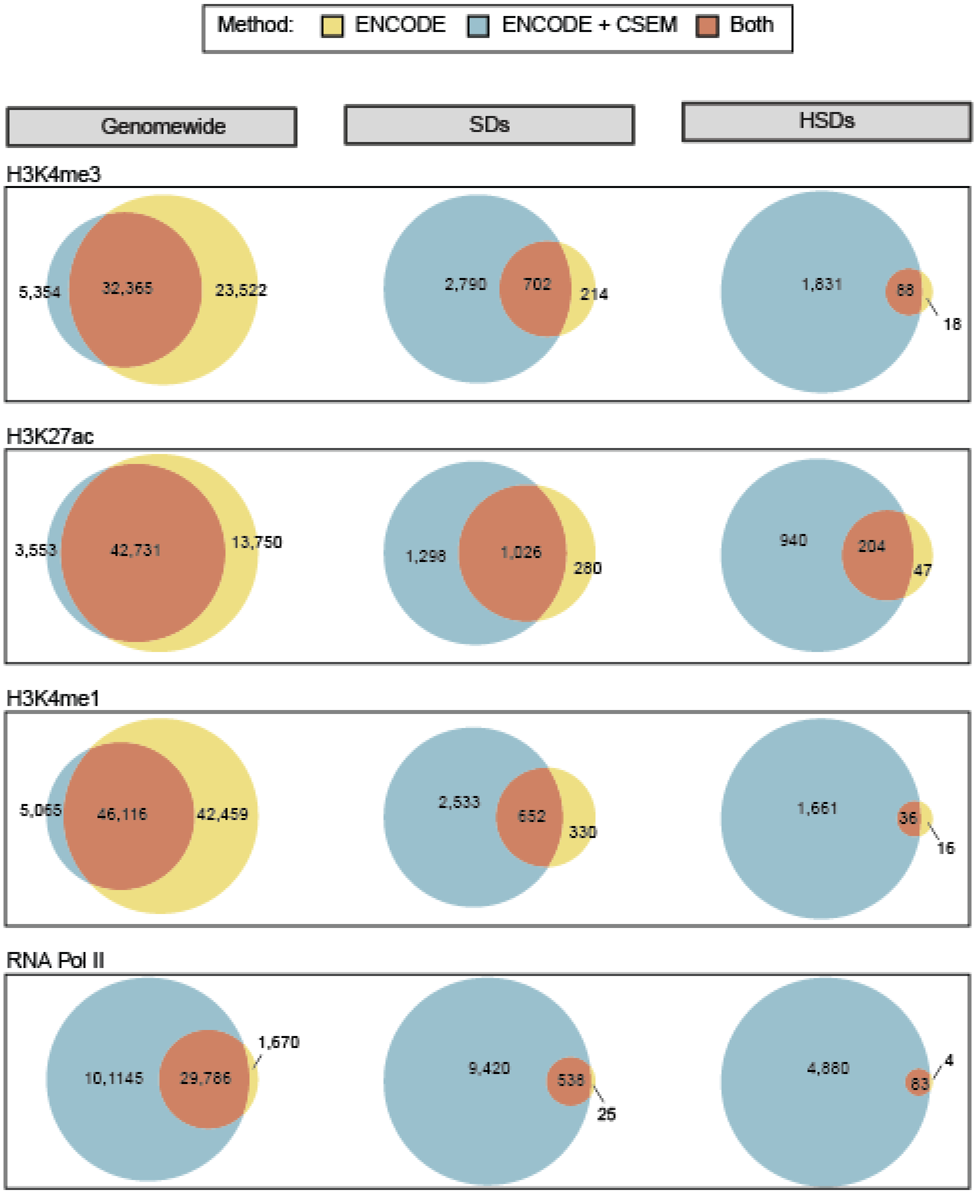


**Figure S5. Comparison of ChIP-seq peaks from ENCODE and ENCODE data with multimapping and CSEM allocation.** Overlap between data sets is shown for the whole genome, SDs, and HSDs (SDs with over 98% identity) for H3K4me1, H3K27ac, H4K4me3, and RNA PolII. Color indicates the method used for read mapping.

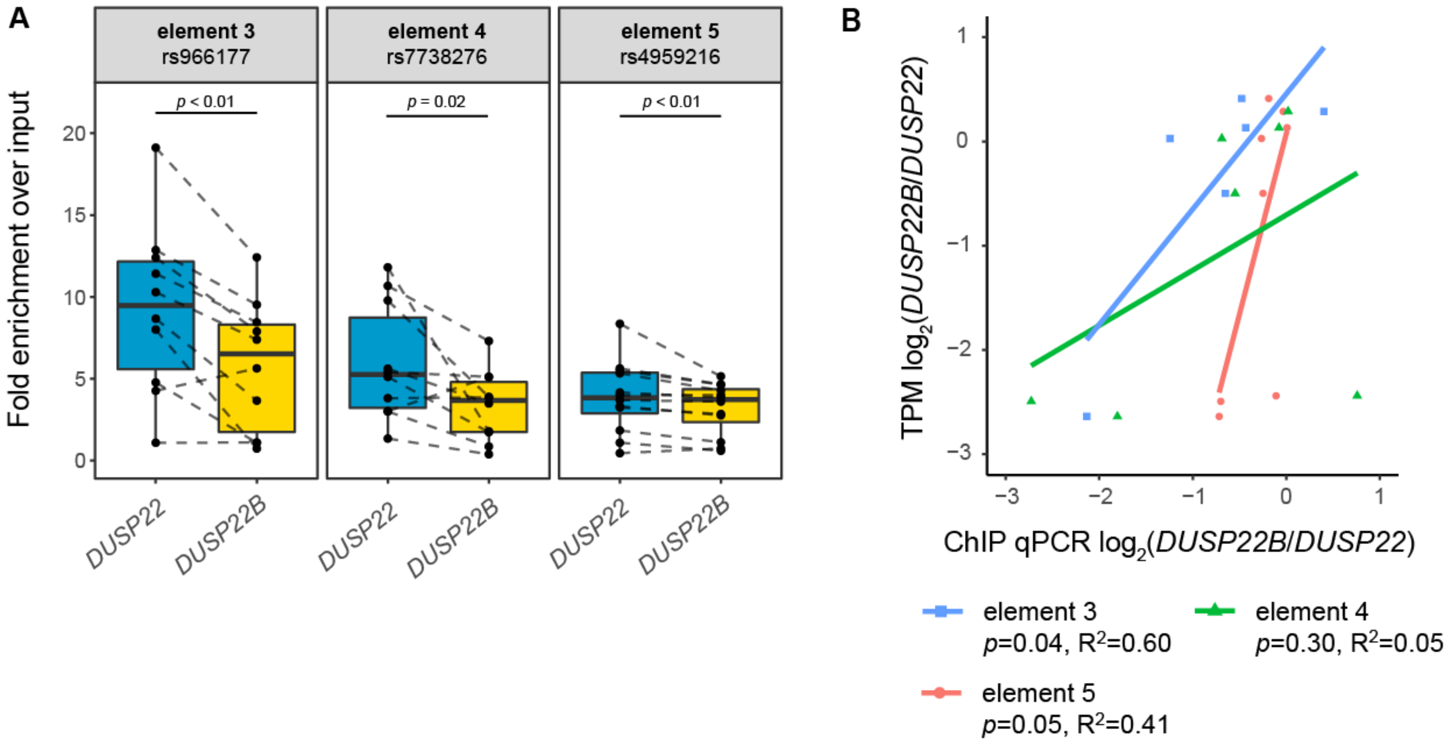


**Figure S6. Paralogous differences of *DUSP22* H3K27ac peaks.** To verify our H3K7ac ChIP-seq results in *DUSP22* and *DUSP22B*, as well as and assess biological reproducibility, we leveraged PSVs (annotated as SNPs in dbSNP) to perform paralog-specific ChIP-qPCR of three H3K27ac-enriched peaks in these genes. **(A)** *DUSP22* and *DUSP22B*-specific enrichment at three putative CREs, with paralogs distinguished using PSVs (N=12, 10, and 10 LCLs, respectively). Each variant lies within an element tested for enhancer activity with a luciferase reporter (Figures 5, S13, S14). Measurements were performed in triplicate and averaged. Differences in enrichment between paralogs were determined with a Wilcoxon signed-rank test, and *p*-values are denoted between the boxplots. **(B)** Correlation of expression divergence ($\log_2$ ratio of TPMs) with differences in enrichment (ChIP-qPCR signal) at the same three variants, for LCLs with RNA-seq data (Pickrell et al. 2010) (N=6, 7, and 8, respectively).

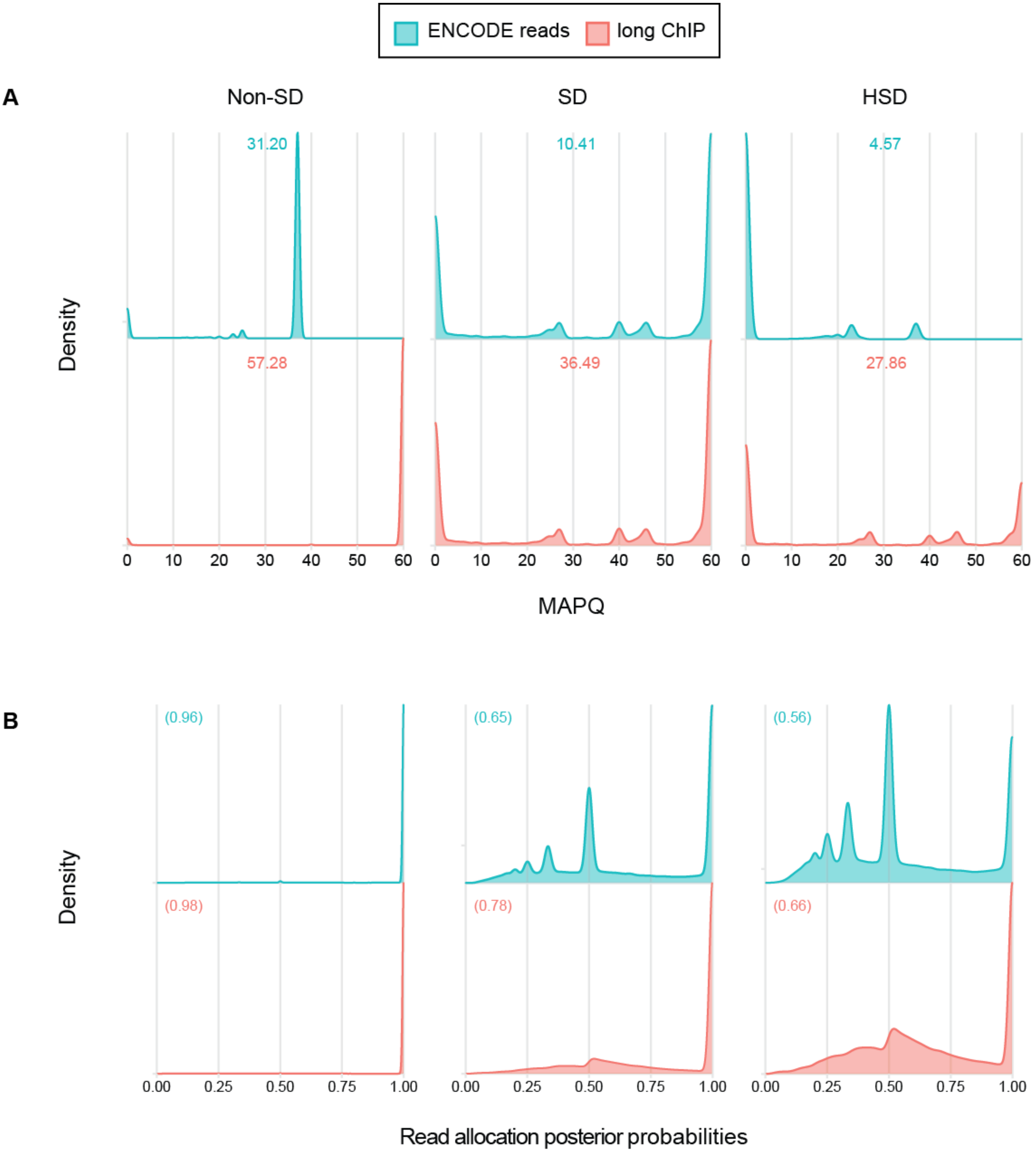


**Figure S7. Summary of long ChIP data analysis (H3K27ac). (A)** Distribution of bwa aln (ENCODE short reads, top) and bwa mem (long ChIP, bottom) alignment mapping quality (MAPQ) scores. Density plots are shown for non-SD regions (left), SDs (middle), and SDs of over 98% sequence identity (HSD, left). The mean MAPQ score is shown on the top of each panel. **(B)** Distribution of bowtie (ENCODE short reads, top) and bowtie2 (long ChIP, bottom) CSEM posterior alignment probabilities. Density plots are shown for the whole genome (left), SDs (middle), and SDs of over 98% sequence identity (HSD, left). The mean posterior probability is shown on the top of each panel.

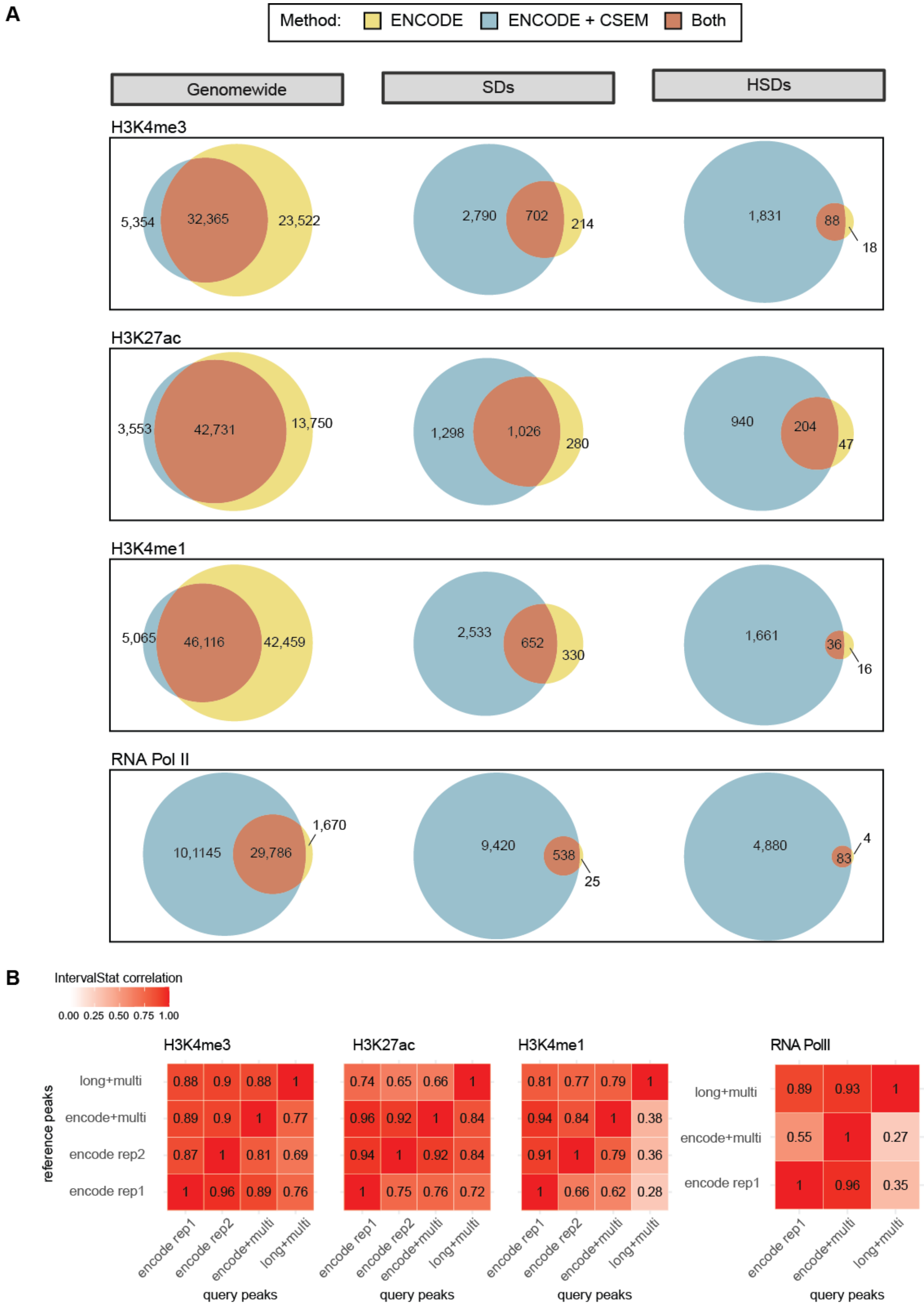


| H3K4me3 | encode rep1 | encode rep2 | encode+multi | long+multi |
|---|---|---|---|---|
| long+multi | 0.88 | 0.9 | 0.88 | 1 |
| encode+multi | 0.89 | 0.9 | 1 | 0.77 |
| encode rep2 | 0.87 | 1 | 0.81 | 0.69 |
| encode rep1 | 1 | 0.96 | 0.89 | 0.76 |

| H3K27ac | encode rep1 | encode rep2 | encode+multi | long+multi |
|---|---|---|---|---|
| long+multi | 0.74 | 0.65 | 0.66 | 1 |
| encode+multi | 0.96 | 0.92 | 1 | 0.84 |
| encode rep2 | 0.94 | 1 | 0.92 | 0.84 |
| encode rep1 | 1 | 0.75 | 0.76 | 0.72 |

| H3K4me1 | encode rep1 | encode rep2 | encode+multi | long+multi |
|---|---|---|---|---|
| long+multi | 0.81 | 0.77 | 0.79 | 1 |
| encode+multi | 0.94 | 0.84 | 1 | 0.38 |
| encode rep2 | 0.91 | 1 | 0.79 | 0.36 |
| encode rep1 | 1 | 0.66 | 0.62 | 0.28 |

| RNA PolII | encode rep1 | encode+multi | long+multi |
|---|---|---|---|
| long+multi | 0.89 | 0.93 | 1 |
| encode+multi | 0.55 | 1 | 0.27 |
| encode rep1 | 1 | 0.96 | 0.35 |

**Figure S8. Comparison of ChIP-seq peaks from long ChIP data with multimapping and CSEM allocation and published ENCODE. (A)** Overlap between data sets is shown for the whole genome, SDs, and HSDs (SDs of over 98% identity) for H3K4me3, H3K27ac, H4K4me1, and RNA PolII. Color indicates the method used for read mapping. **(B)** Pairwise correlations of genome-wide peak sets from single ENCODE replicates, ENCODE multi-mapping with CSEM allocation, and long ChIP multimapping with CSEM allocation. Unidirectional correlations were determined by IntervalStats (Chikina and Troyanskaya 2012), with overlapping peaks defined at $p<0.05$.

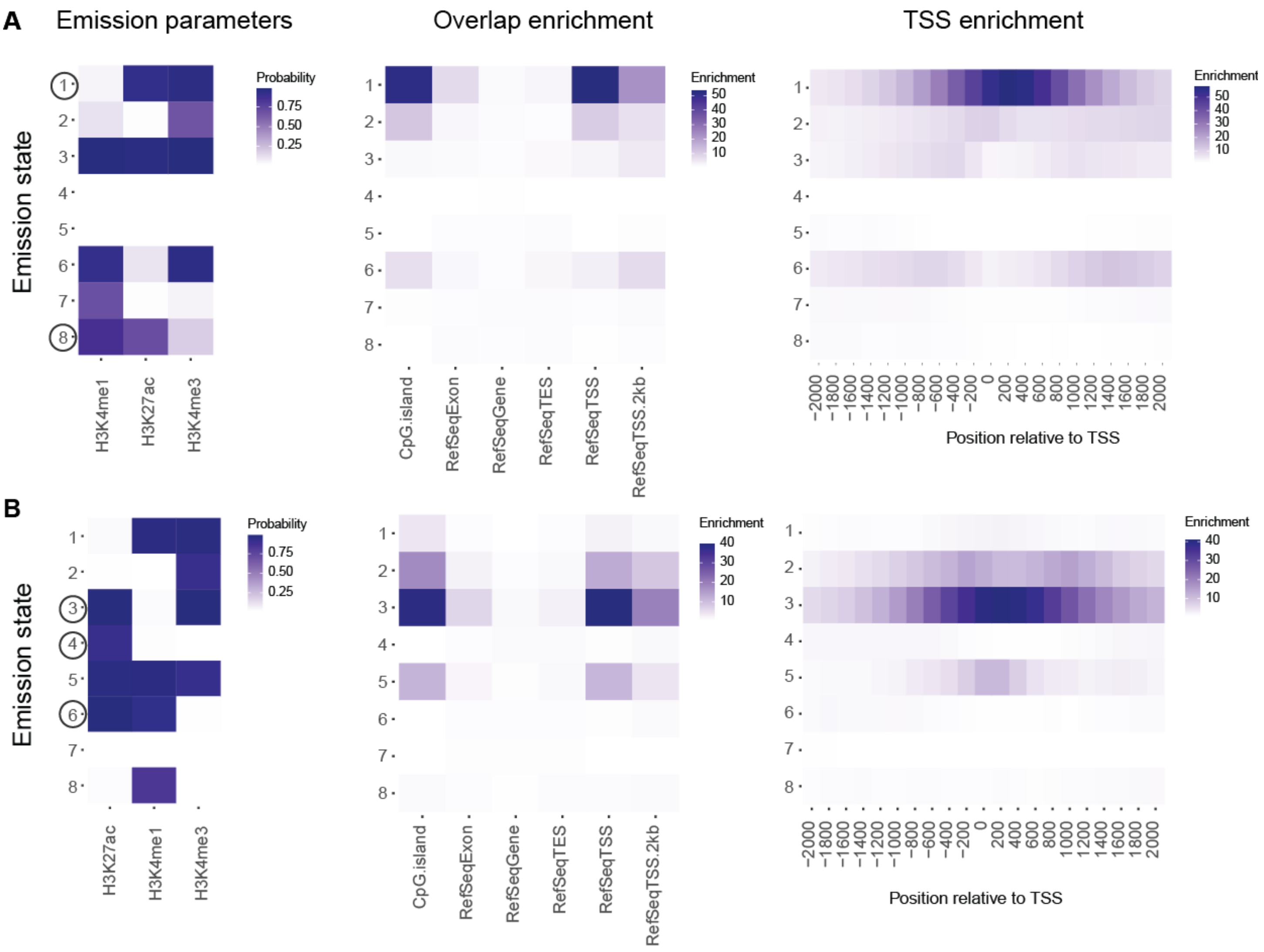


**Figure S9. ChromHMM models used to cCREs. (A)** Emission parameters, genomic features overlap, and enrichment relative to transcription start sites (TSSs) of an 8-state ChromHMM model built on ENCODE data (multiple-mapping and CSEM allocation). Darker blue indicates a higher probability of observing a given histone mark in each state, or a higher fold-enrichment in a given genomic feature or distance from TSS. State 1 was chosen to represent active promoters, and state 8 was chosen for active enhancers (circled). **(B)** Emission parameters, genomic features overlap, and enrichment relative to TSSs for an 8-state ChromHMM model built on long ChIP data (multiple-mapping and CSEM allocation). State 3 was chosen for active promoters, and states 4 and 6 were chosen for active enhancers (circled).

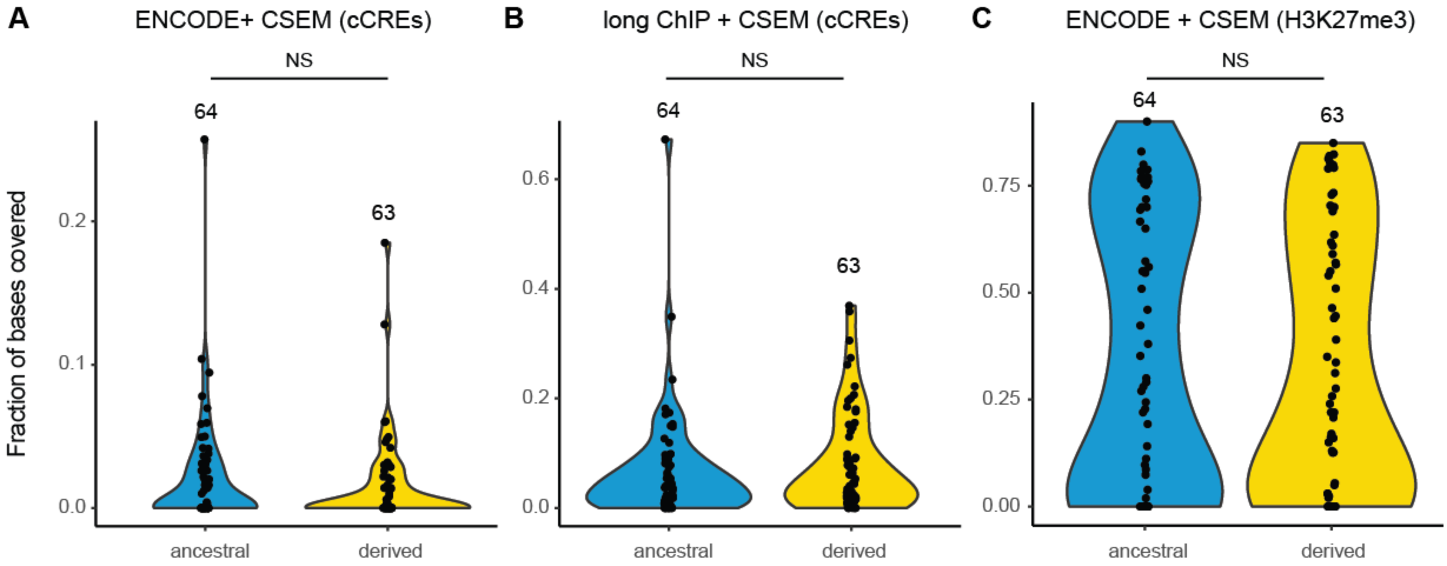


**Figure S10. Global comparison of ancestral and derived HSDs.** Violin plots represent the fraction of bases covered by (A) ENCODE multi-mapping cCREs, (B) long ChIP multi-mapping cCREs, and (C) ENCODE multi-mapping H3K27me3 domains. Fractional coverage was calculated in 100-kb windows for ancestral and derived HSD regions. Values were compared with a Wilcoxon signed-rank test.

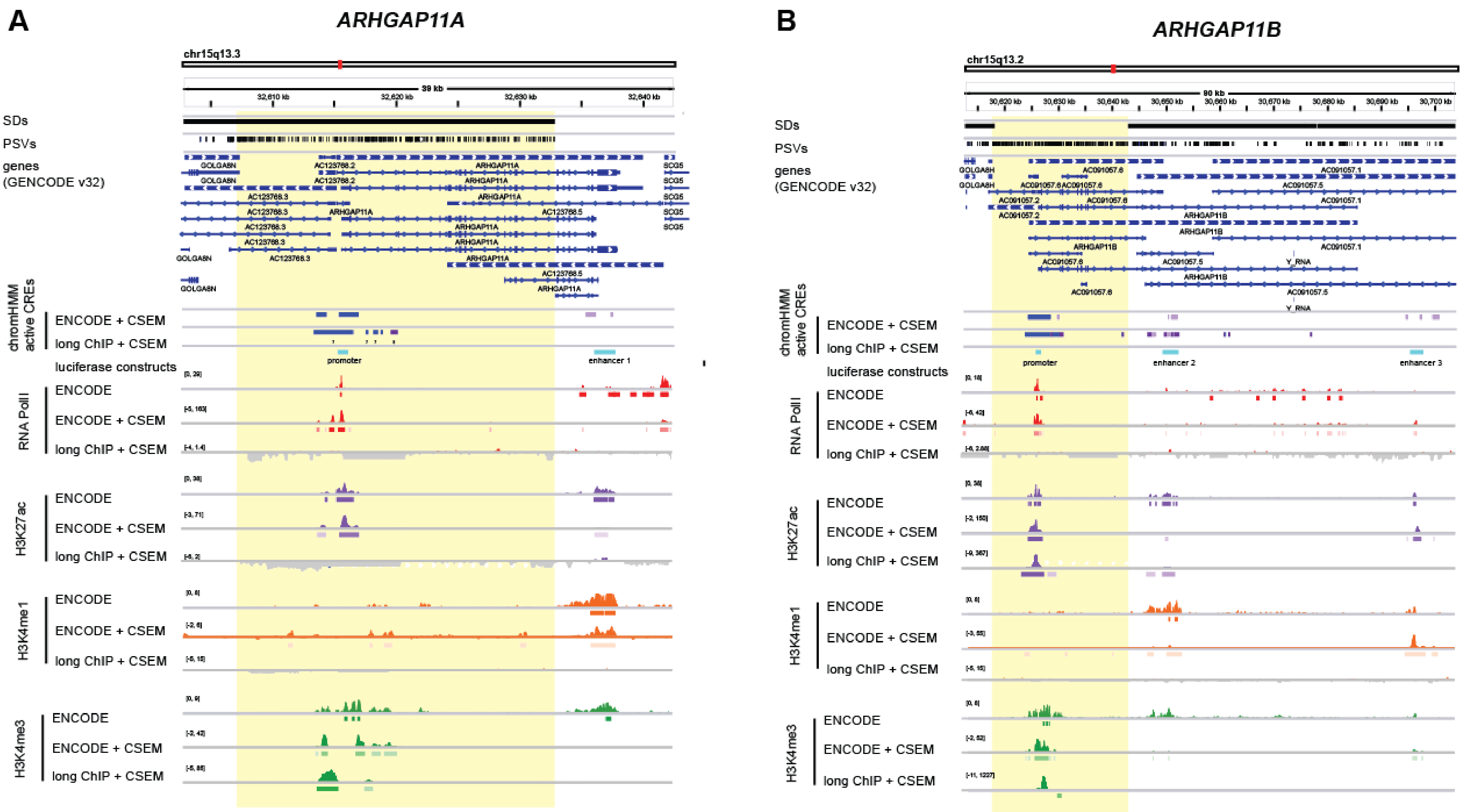


**Figure S11. Epigenetic landscape of *ARHGAP11* genes. (A)** *ARHGAP11A*. **(B)** *ARHGAP11B*. Coordinates indicate location on chromosome 15. The paralogous duplicated region is highlighted in yellow. Segmental duplications (SDs) and paralog-specific variants (PSVs) are indicated with black bars. ChromHMM segmentations are shown for active promoters (blue) and enhancers (lavender), as defined on ENCODE and long ChIP data (multimapping with CSEM allocation). Regions cloned and tested with luciferase reporters are shown in cyan. For each ChIP-seq target, a signal track is shown for published ENCODE; reanalyzed, multimapped ENCODE with CSEM allocation; and multimapped long ChIP with CSEM allocation. Visualized with the Integrative Genomics Viewer.

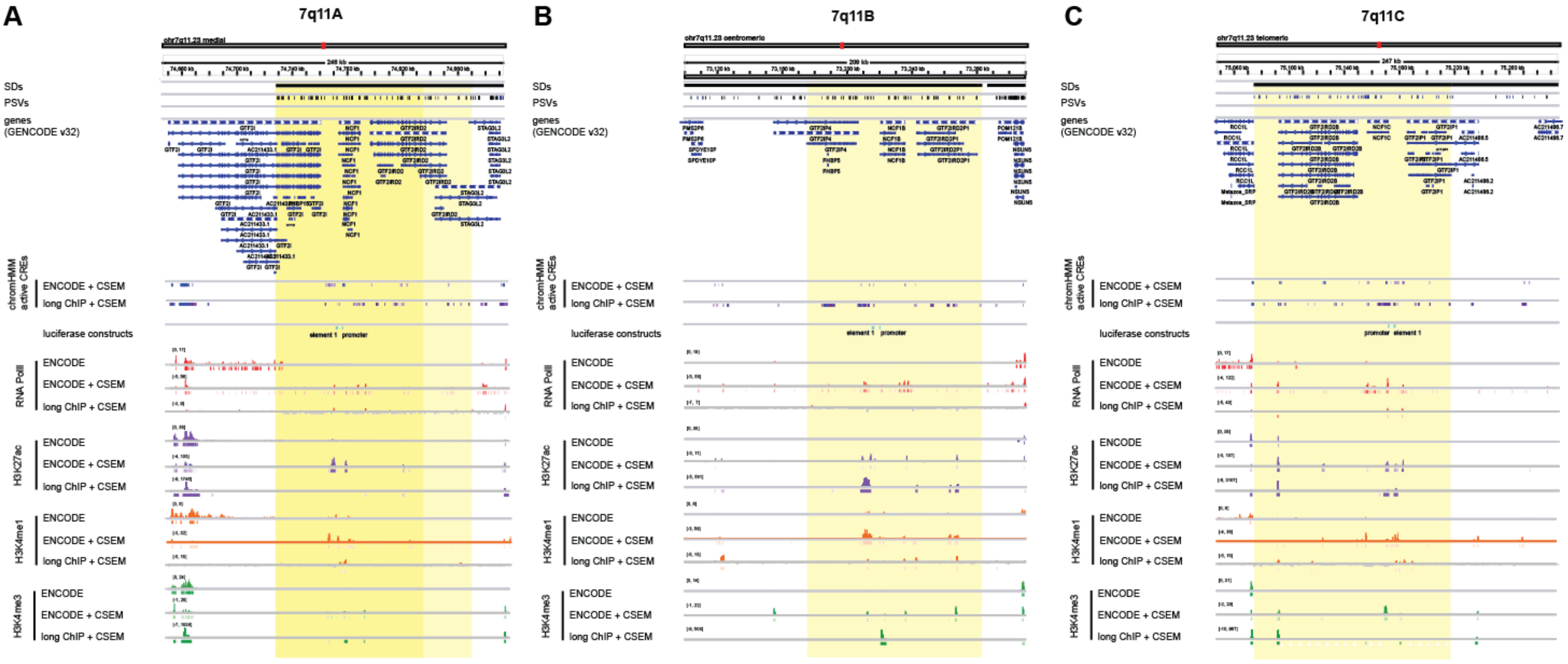


**Figure S12. Epigenetic landscape of chromosome 7q11. (A)** Ancestral locus. **(B)** Primary derived (centromeric) locus. **(C)** Secondary derived (telomeric) locus. Coordinates indicate location on chromosome 7. The paralogous duplicated regions are highlighted in yellow. Segmental duplications (SDs) and paralog-specific variants (PSVs) are indicated with black bars ChromHMM segmentations are shown for active promoters (blue) and enhancers (lavender), as defined on ENCODE and long ChIP data (multimapping with CSEM allocation). Regions cloned and tested with luciferase reporters are shown in cyan. For each ChIP-seq target, a signal track is shown for published ENCODE; reanalyzed, multimapped ENCODE with CSEM allocation; and multimapped long ChIP with CSEM allocation. Visualized with the Integrative Genomics Viewer.

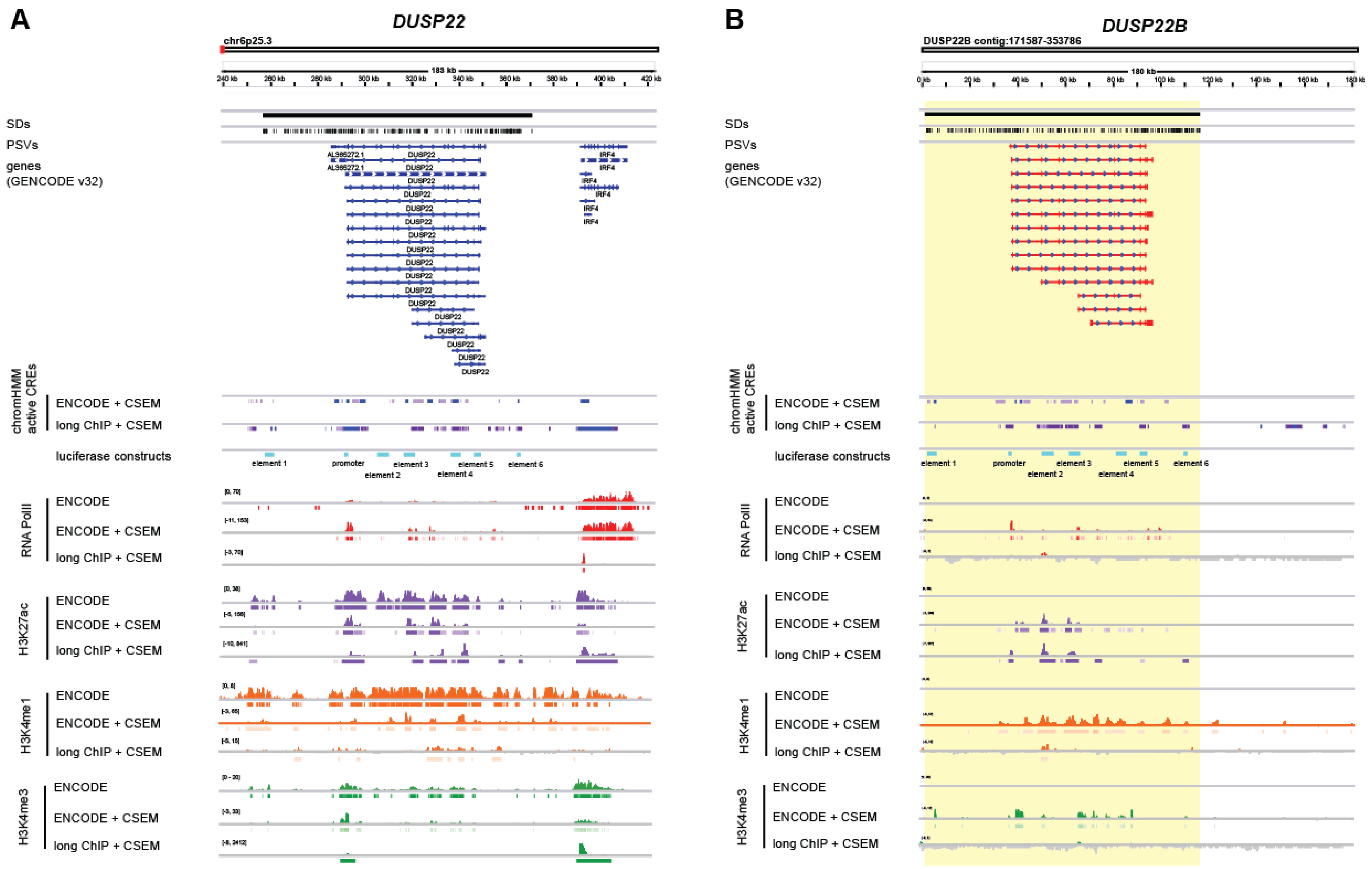


**Figure S13. Epigenetic landscape of *DUSP22* genes. (A)** *DUSP22*. **(B)** *DUSP22B*. Coordinates indicate location on chromosome 6 or the *DUSP22B* contig as published in (Dennis et al. 2017). *DUSP22B* gene models depict BLAT alignments of *DUSP22* transcripts. The paralogous duplicated region is highlighted in yellow. Segmental duplications (SDs) and paralog-specific variants (PSVs) are indicated with black bars. ChromHMM segmentations are shown for active promoters (blue) and enhancers (lavender), as defined on ENCODE and long ChIP data (multimapping with CSEM allocation). Regions cloned and tested with luciferase reporters are shown in cyan. For each ChIP-seq target, a signal track is shown for published ENCODE; reanalyzed, multimapped ENCODE with CSEM allocation; and multimapped long ChIP with CSEM allocation. Visualized with the Integrative Genomics Viewer.

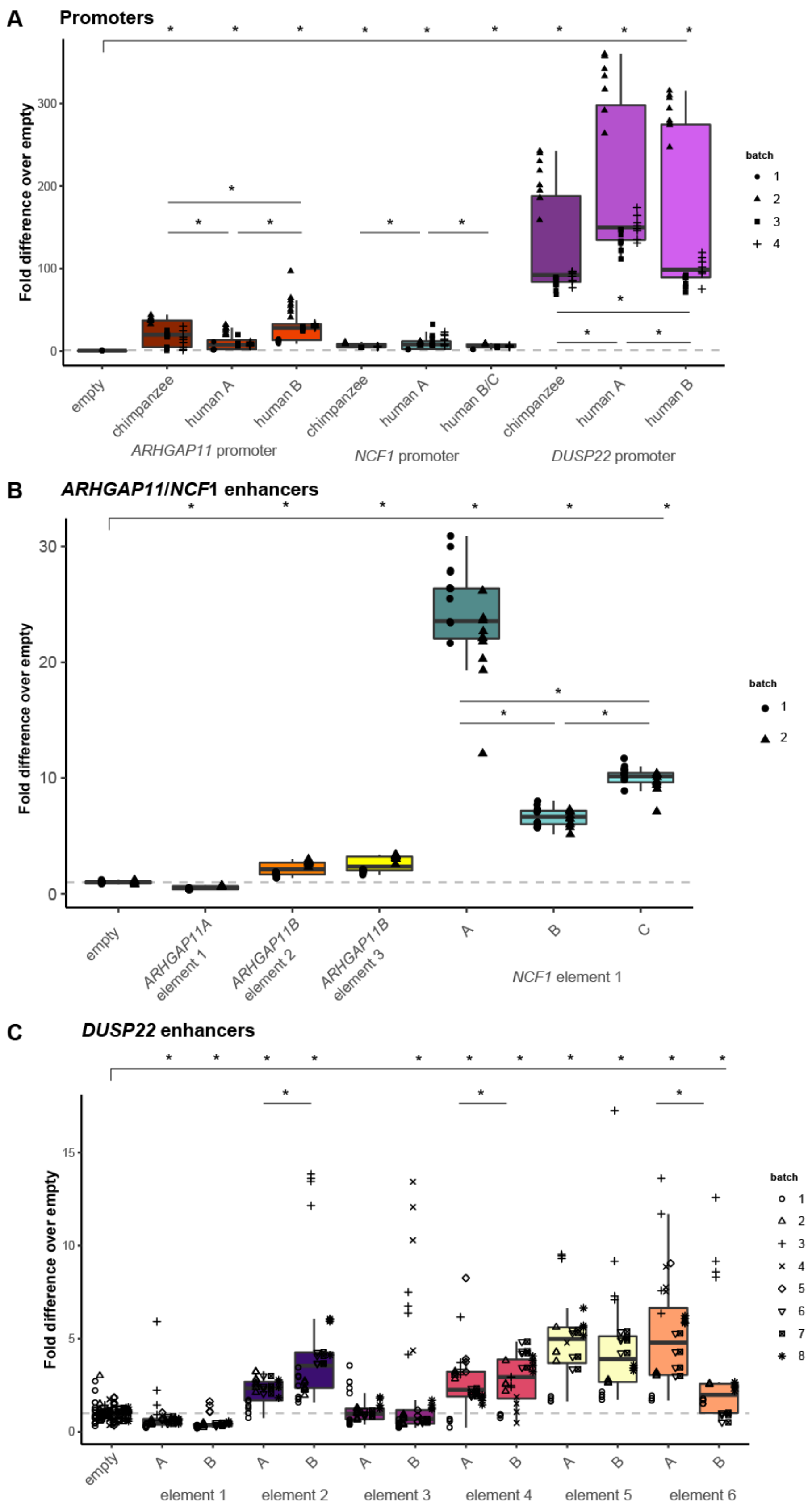


**Figure S14 Luciferase activity of candidate CREs from all HeLa experiments.** Luciferase activity for each cloned construct is shown as the fold difference over the average negative control value for **(A)** promoters, **(B)** *ARHGAP11* and *NCF1* candidate enhancers, and **(C)** *DUSP22* candidate enhancers. Values are visually separated by experimental batch. Significant differences ($p < 0.05$, Tukey post-hoc test of two-way ANOVA) from empty (top bar) and between homologous sequences are indicated with an asterisk.

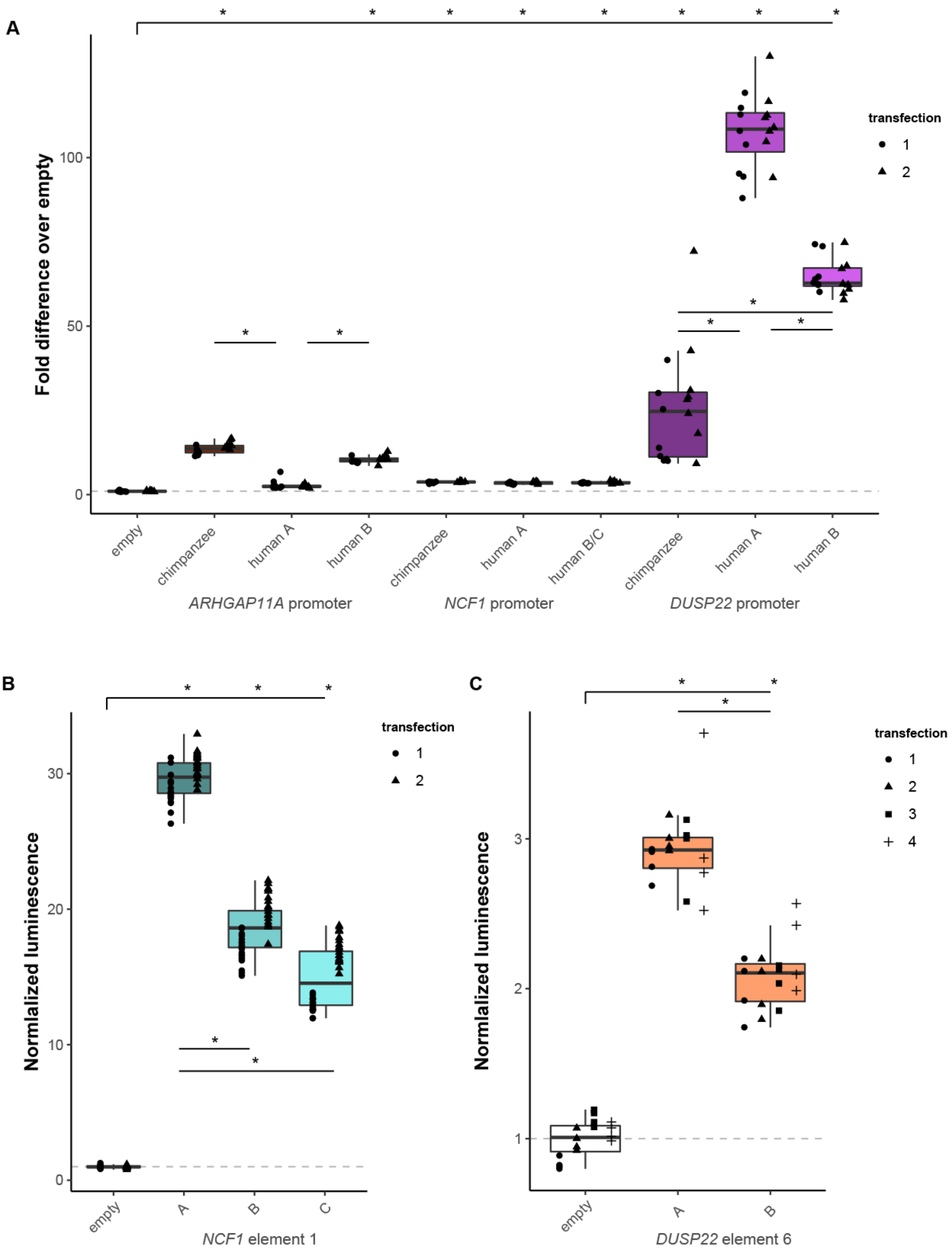


**Figure S15. Luciferase activity of candidate CREs from all LCL experiments.** Luciferase activity for each cloned construct is shown as the fold difference over the average negative control value for **(A)** promoters, **(B)** *NCF1* candidate element 1, and **(C)** *DUSP22* candidate element 6. Values are visually separated by experimental batch. Significant differences ($p < 0.05$, Tukey post-hoc test of two-way ANOVA) from empty (top bar) and between homologous sequences are indicated with an asterisk.

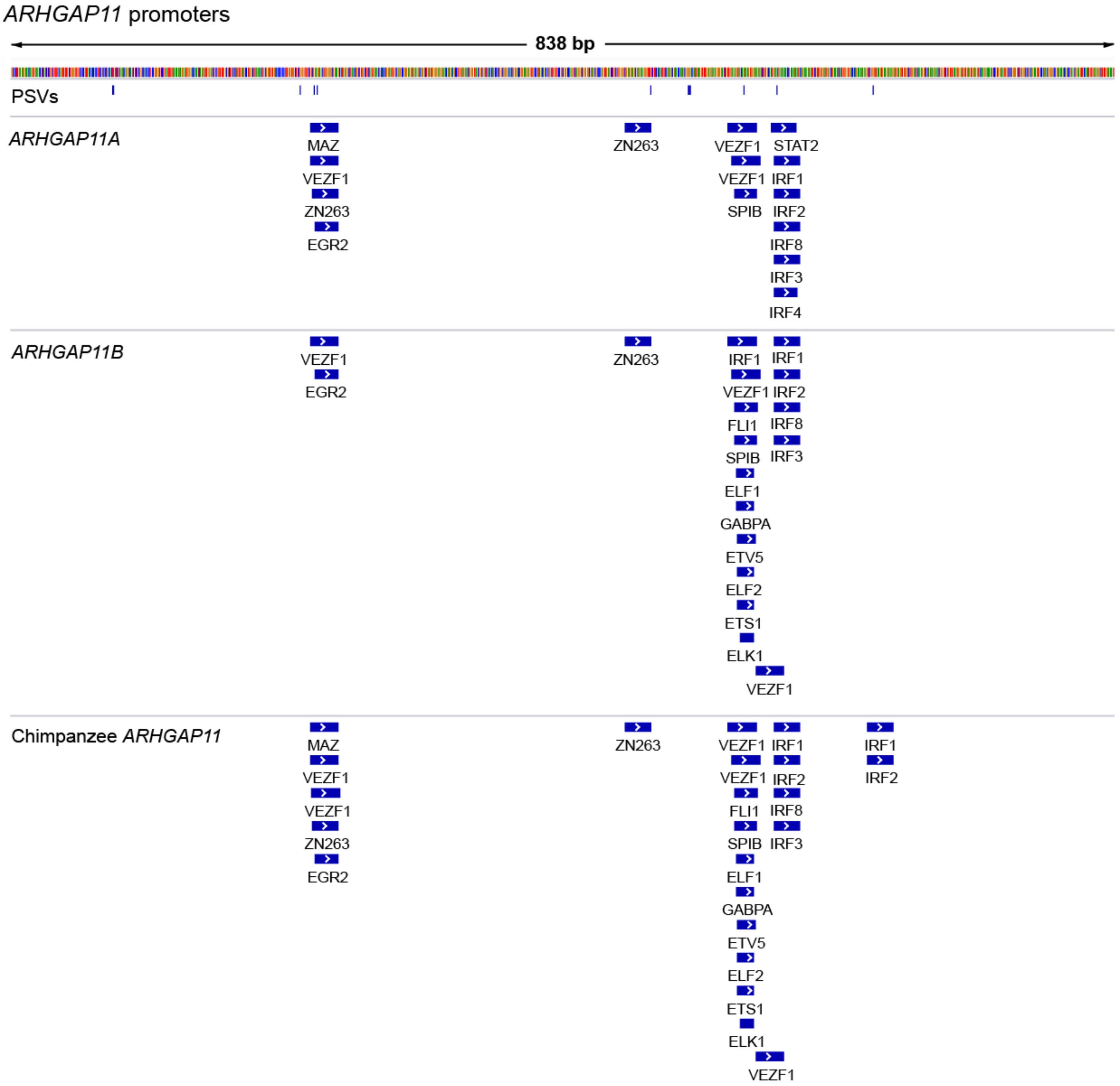


**Figure S16. Transcription factor binding sites (TFBSs) identified in *ARHGAP11* promoters.** Significant matches (5% FDR) for HOCOMOCO v.12 TFBS motifs intersecting PSVs are depicted under the cloned sequence tested with luciferase reporter. PSVs are depicted with blue vertical lines. Visualized with the Integrative Genomics Viewer.

### References

Chikina MD, Troyanskaya OG. 2012. An effective statistical evaluation of ChIPseq dataset similarity. *Bioinformatics* 28:607–613.

Dennis MY, Harshman L, Nelson BJ, Penn O, Cantsilieris S, Huddleston J, Antonacci F, Penewit K, Denman L, Raja A, et al. 2017. The evolution and population diversity of human-specific segmental duplications. *Nat Ecol Evol* 1:69.

Nguyen LS, Jolly L, Shoubridge C, Chan WK, Huang L, Laumonnier F, Raynaud M, Hackett A, Field M, Rodriguez J, et al. 2012. Transcriptome profiling of UPF3B/NMD-deficient lymphoblastoid cells from patients with various forms of intellectual disability. *Mol. Psychiatry* 17:1103–1115.

Pickrell JK, Marioni JC, Pai AA, Degner JF, Engelhardt BE, Nkadori E, Veyrieras J-B, Stephens M, Gilad Y, Pritchard JK. 2010. Understanding mechanisms underlying human gene expression variation with RNA sequencing. *Nature* [Internet] 464:768–772. Available from: http://dx.doi.org/10.1038/nature08872
